# Deep divergence and genomic diversification of gut symbionts of neotropical stingless bees

**DOI:** 10.1101/2022.12.08.519137

**Authors:** Garance Sarton-Lohéac, Carlos Gustavo Nunes da Silva, Florent Mazel, Gilles Baud, Vincent de Bakker, Sudip Das, Yassine El Chazli, Kirsten Ellegaard, Marc Garcia-Garcera, Natasha Glover, Joanito Liberti, Lorena Nacif Marçal, Aiswarya Prasad, Vincent Somerville, SAGE class 2019-2020 and 2020-2021, Germán Bonilla-Rosso, Philipp Engel

## Abstract

Social bees harbor conserved gut microbiota that may have been acquired in a common ancestor of social bees and subsequently co-diversified with their hosts. However, most of this knowledge is based on studies on the gut microbiota of honey bees and bumble bees. Much less is known about the gut microbiota of the third and most diverse group of social bees, the stingless bees. Specifically, the absence of genomic data from their microbiota presents an important knowledge gap in understanding the evolution and functional diversity of the social bee microbiota. Here we combined community profiling with culturing and genome sequencing of gut bacteria from six neotropical stingless bee species from Brazil. Phylogenomic analyses show that most stingless bee gut isolates form deep-branching sister clades of core members of the honey bee and bumble bee gut microbiota with conserved functional capabilities, confirming the common ancestry and ecology of their microbiota. However, our bacterial phylogenies were not congruent with those of the host indicating that the evolution of the social bee gut microbiota was not driven by strict co-diversification, but included host switches and independent symbiont gain and losses. Finally, as reported for the honey bee and bumble bee microbiota, we find substantial genomic divergence among strains of stingless bee gut bacteria suggesting adaptation to different host species and glycan niches. Our study offers first insights into the genomic diversity of the stingless bee microbiota, and highlights the need for broader samplings to understand the evolution of the social bee gut microbiota.

**Importance:** Stingless bees are the most diverse group of the corbiculate bees and represent important pollinator species throughout the tropics and subtropics. They harbor specialized microbial communities in their gut that are related to those found in honey bees and bumble bees and that are likely important for bee health. Few bacteria have been cultured from the gut of stingless bees which has prevented characterization of their genomic diversity and functional potential. Here, we established cultures of major community members of the gut microbiota of six stingless bee species and sequenced their genomes. We find that most stingless bee isolates belong to novel bacterial species distantly related to those found in honey bees and bumble bees and encoding similar functional capabilities. Our study offers a new perspective on the evolution of the social bee gut microbiota and presents the basis to characterize the symbiotic relationships between gut bacteria and stingless bees.

## Introduction

The eusocial corbiculate bees (hereafter social bees) comprise more than 700 species distributed in three distinct tribes: honey bees (Apini), bumble bees (Bombini), and stingless bees (Meliponini). Stingless bees and bumble bees form a monophyletic clade, which is sister to the honey bees, with whom they shared a common ancestor 80-100 million years ago (1–3). Like mammals, social bees harbor dense and specialized bacterial communities in their gut that affect bee health and behavior (4–12). The composition of the gut microbiota of social bees is relatively simple typically consisting of <10 bacterial phylotypes, i.e. sequence clusters sharing >97% identity in the 16S rRNA gene (13–20). Five of these phylotypes (*Snodgrassella*, *Gilliamella*, Bombilactobacillus Firm-4, *Lactobacillus* Firm-5, and *Bifidobacterium*) have been referred to as the core gut microbiota of the social bees (20), because they are prevalent and abundant across honey bee, bumble bee, and stingless bees. Most members of the bee gut microbiota are culturable and gnotobiotic bees can be generated for several species (5, 21–23). Together, these distinctive characteristics make the social bee microbiota a versatile model system for studying the evolution and ecology of host-associated microbial communities. Moreover, social bees are important pollinators that suffer from severe population declines (24, 25) which makes studies of their microbiota relevant in its own right.

Most of what is currently known about the gut microbiota of social bees stems from studies on honey bees and bumble bees. Genomic and experimental approaches have revealed that their gut bacteria are usually saccharolytic fermenters that utilize plant glycans derived from the pollen and nectar/honey diet of the host (22, 23, 26–32). Further, it has been shown that the core members of the honey bee and bumble bee gut microbiota have substantially diversified (27–30, 33–36). They consist of divergent sub-lineages (or species) and contain large extents of strain-level diversity and gene content variation. Most sub-lineages are host-specific (23, 27, 33, 36), and their phylogenetic relationships are to some extent congruent with the phylogeny of the host (16, 33). Therefore, it has been suggested that the core members of the microbiota were acquired in a common ancestor of the social bees (20) and possibly co-diversified with the host (16, 33). In addition, studies in the Western honey bee (*Apis mellifera*) have shown that the diversification of the bee gut microbiota was also driven by adaptation to different spatial and metabolic niches within the gut (27, 30, 33–35). For example, strains of closely related sub-lineages of *Lactobacillus* Firm-5 and *Bifidobacterium* can coexist in individual bees. They carry distinct gene sets for the breakdown and utilization of pollen-derived carbohydrates which allows them to partition the available dietary glycan niches in the gut (27, 30, 34).

In contrast to the microbiota of honey bees and bumble bees, much less is known about the gut microbiota of the third group of social bees, the stingless bees (Meliponini). Previous studies have focused on determining the taxonomic composition of the gut microbiota of these bees using 16S rRNA gene sequencing (15, 17, 20, 37–39). However, only a few bacteria have been cultured from stingless bees (40–42) and, except for two strains of *Bombilactobacillus* Firm-4 recently isolated from bees from Australia (43), no genomic data is currently available for core members of the gut microbiota of stingless bees.

With >500 described species, stingless bees present the largest and most diverse group of the social bees (44, 45). They are naturally distributed throughout the tropical and subtropical regions of Africa, Asia, Australia, and the Americas and exhibit great variation in morphology, diet, foraging range, social structure, and nesting habits (44, 45). As host phylogeny and ecology are both key determinants of gut microbiota composition (46–51), we hypothesize that genomic studies on bacterial isolates will help us to understand the functional diversity of gut bacteria of stingless bees and provide novel insights into the evolution of these bacteria across social bees, specifically in respect of the possible co-diversification with the host.

To address these questions, we looked at the gut microbiota of six neotropical species of stingless bees from Brazil: *Frieseomelitta varia* (Fv), *Scaptotrigona polysticta* (Sp), *Melipona fuliginosa* (Mf), *Melipona interrupta* (Mi), *Melipona seminigra* (Ms), and *Melipona lateralis* (Ml). We determined the composition of the gut microbiota of these bees using 16S rRNA gene sequencing, established a comprehensive culture collection of bacterial isolates, and conducted genome sequencing and comparative genomics to determine the phylogenetic placement, genomic diversity, and functional capabilities of these bacteria relative to those previously isolated from honey bees and bumble bees.

## Results

### Six neotropical stingless bee species from Brazil harbor distinct gut microbiota dominated by nine bacterial families

We sampled three colonies of six stingless bee species (Fv, Sp, Mf, Mi, Ms, Ml) from a meliponary located in the Amazonian rainforest near Manaus (**Supplementary Table 1**). For each colony, we pooled the guts of 15-60 worker bees (depending on the size of the bee species, see Methods) before DNA isolation. The V4 region of the 16S rRNA gene was amplified and sequenced with Illumina MiSeq 2×250bp resulting in a median depth of 64,713 (52,479 to 95,774) reads per sample. In total, we identified 277 amplicon sequence variants (ASVs, 29-63 ASVs per sample), which belonged to 36 different bacterial families (**Supplementary Table 2**). Despite this diversity, only nine families dominated the samples, representing together 97% of all quality-filtered reads (93-99% of the reads per sample): Acetobacteraceae, Bifidobacteriaceae, Enterobacteriaceae, Lactobacillaceae, Neisseriaceae, Orbaceae, Prevotellaceae, Streptococcaceae and Veillonellaceae (**Figure 1A**).

**Figure 1.**
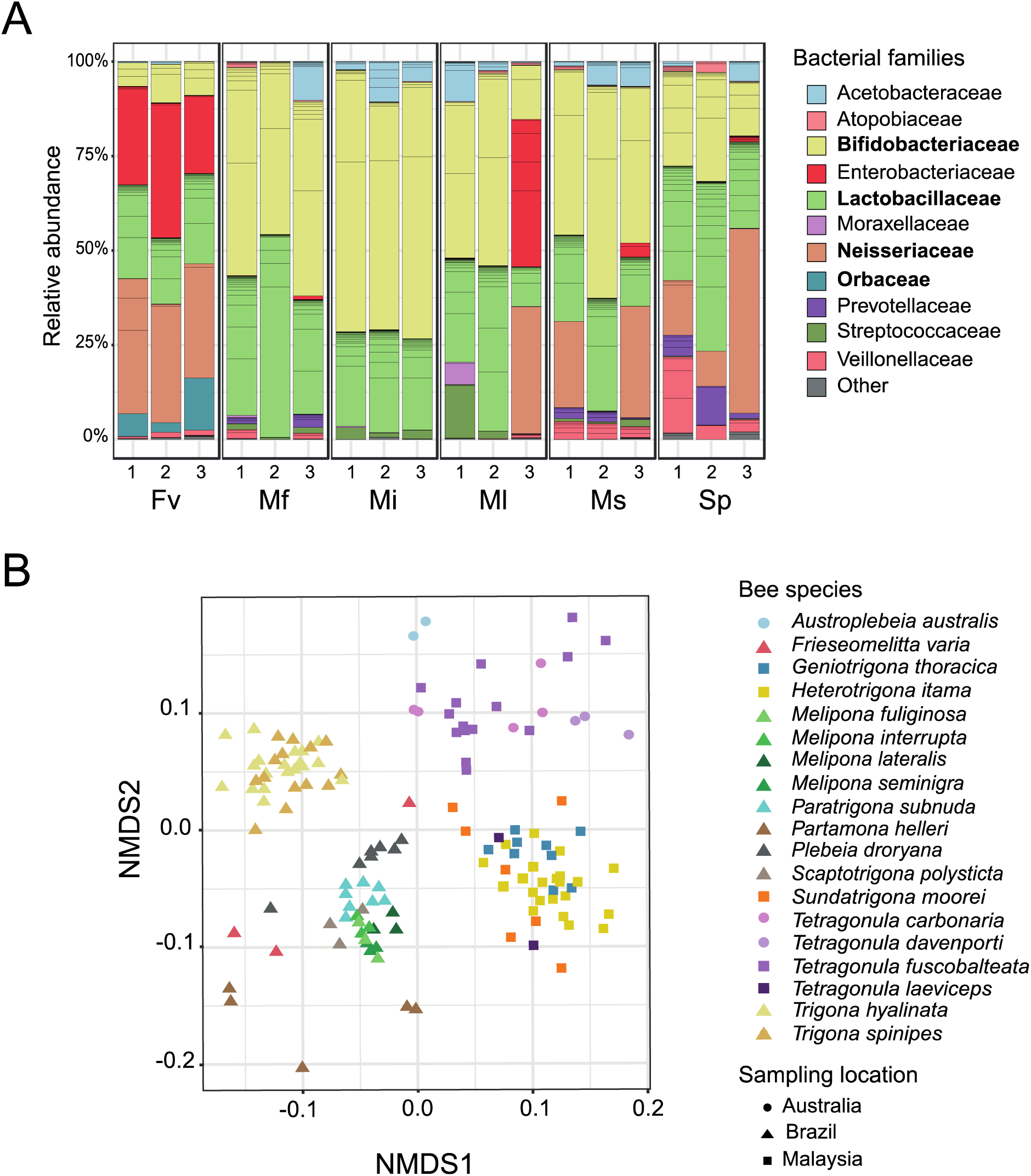
Community analysis of the gut microbiota of stingless bees. **(A)** 16S rRNA gene-based community profiles of the gut microbiota of three colonies of six different stingless bee species collected in Brazil. Fv – *Frieseomelitta varia,* Ms – *Melipona seminigra*, Ml – *Melipona lateralis*, Mf – *Melipona fuliginosa*, Mi – *Melipona interrupta*, Sp – *Scaptotrigona polysticta*. Relative abundance of ASVs is shown. ASVs are ordered and colored at the family level according to the legend, families of core members are displayed in bold. ASVs with <1% relative abundances are summed up as ‘others’ and shown in grey. **(B)** NMDS based on ASV relative abundance (Bray-Curtis dissimilarity) across 136 16S rRNA gene amplicon sequence samples including data from a previous study (20) and our study.

While Lactobacillaceae and Bifidobacteriaceae were abundant across all samples, there were clear differences in the distribution of some of the other bacterial families (**Figure 1A**). Neisseriaceae were abundant in the samples of Sp and Fv, but were only detected in three out of 12 samples from the genus *Melipona* (**Figure 1A, Supplementary Table 2**). In contrast, Acetobacteraceae and Streptococcaceae were present in most *Melipona* samples but rare across samples of Fv and Sp. Orbaceae and Enterobacteriaceae were mostly detected in the three Fv samples. Intriguingly, a single Enterobacteriaceae ASV constituted the most abundant community member in this bee species (21 – 36% of the reads per sample). According to these compositional differences, non-metric multidimensional scaling (NMDS) separated the samples into three distinct clusters: two clusters comprised all samples of Fv and Sp, respectively, and the third cluster comprised all samples of the four *Melipona* species (Mf, Mi, Ms, Ml) (**Supplementary Figure 1A**).

### Related stingless bee species have overlapping community profiles

We compared our results to a previously published amplicon sequencing dataset from stingless bees (20) to assess the similarity of the communities to those of other stingless bee species. After discarding samples with <5,000 reads to control for variation in sequencing depths, our dataset comprised 135 samples from 19 different host species and three different countries. We detected 688 ASVs in total with a median of 18 ASVs per sample (3 to 63 ASVs) (**Supplementary Table 3**), spanning 53 bacterial families. Overall, the taxonomic patterns were similar across the analyzed bee species. Apart from one sample of *Tetragonula fuscobalteata,* for which 99% of the reads belonged to a single Weeksellaceae ASV, the nine families dominating in the six bee species from our study were also abundant in the microbiota of the samples from the previous study and represented between 34% to 100% of the total number of reads (**Supplementary Figure 1B, Supplementary Table 3**).

NMDS based on ASV relative abundances separated the samples by location (i.e. samples from Brazil were different from those from Australia and Malaysia, Permutational multivariate analysis of variance (PERMANOVA)**: ‘**Location’ pseudo-F: 20.17, p-value=0.001), and by bee genus (PERMANOVA: ‘Genus’ pseudo-F=10.49, p-value=0.001), although taxonomy and geography are not independent (**Figure 1B**). In contrast, there was only weak clustering at the species level (‘Species’ pseudo-F=1.96, p-value=0.03). Notably, only 30% of all ASVs (206 of 688 ASVs) were shared across host species (i.e. 70% of all ASVs are only found in one species), and most of them (83.4%) only between 2-5 species (**Supplementary Figure 1C**). However, the shared ASVs belonged to the nine predominant bacterial families and represented a large fraction of the total number of reads per sample (53.3-99.7 %; except for the samples of Fv and *P. helleri*: 10.6-15 % of the reads) (**Supplementary Figure 1D**). In particular, bees sampled in the same country or belonging to the same bee genus shared the same ASVs, explaining the clustering of these samples in the NMDS analysis. Together, these results show that despite the large variability observed, the gut microbiota of most stingless bee species is dominated by a few bacterial families and that bee species of the same genus, or with overlapping geographic distribution, have similar community profiles at the 16S rRNA gene level.

### Establishment of a strain collection of gut bacteria isolated from stingless bees

To enable genomic and experimental analyses of stingless bee gut bacteria, we established a culture collection of bacteria isolated from Fv, Sp, Mf, Mi, Ms, and Ml. We plated homogenized gut samples of the six bee species on eight different semi-solid media and under three different atmospheres (microaerobic and anaerobic). This resulted in the cultivation of 98 distinct bacterial isolates (i.e. different 16S rRNA genotype or isolated from a different bee species or colony) from 11 bacterial families (**Figure 2A, Supplementary Table 4**). Most bacteria grew under both microaerobic or anaerobic conditions on generic growth media and formed colonies after 2-4 days of growth. The 16S rRNA genotypes of the isolated strains matched to 32 ASVs, accounting for 16 – 87 % of the overall community of the six stingless bee species and including many shared ASVs (**Figure 2A** and **B**). BLASTN searches of the 16S rRNA gene sequences revealed that many of the isolates (55/98) were related to bacterial strains obtained from the gut of honey bees and bumble bees (such as *Lactobacillus apis*, *Bifidobacterium commune*, *Gilliamella sp.,* or *Snodgrassella alvi*) suggesting that they represent stingless bee isolates of core members of the social bee microbiota. Other isolates had best BLASTN hits to bacteria from other environments, such as *Floricoccus* sp. Isolated from flowers, various Enterobacteriaceae (e.g. *Pantoea* sp., *Klebsiella* sp., *Rosenbergiella epipactidis*) isolated from humans and water, or *Fructobacillus* isolated from flowers and fruits. The percent identity of many of the blast hits was relatively low (<98%) suggesting that the isolated strains potentially correspond to new bacterial species (**Figure 2A** and **C, Supplementary Table 4**).

**Figure 2.**
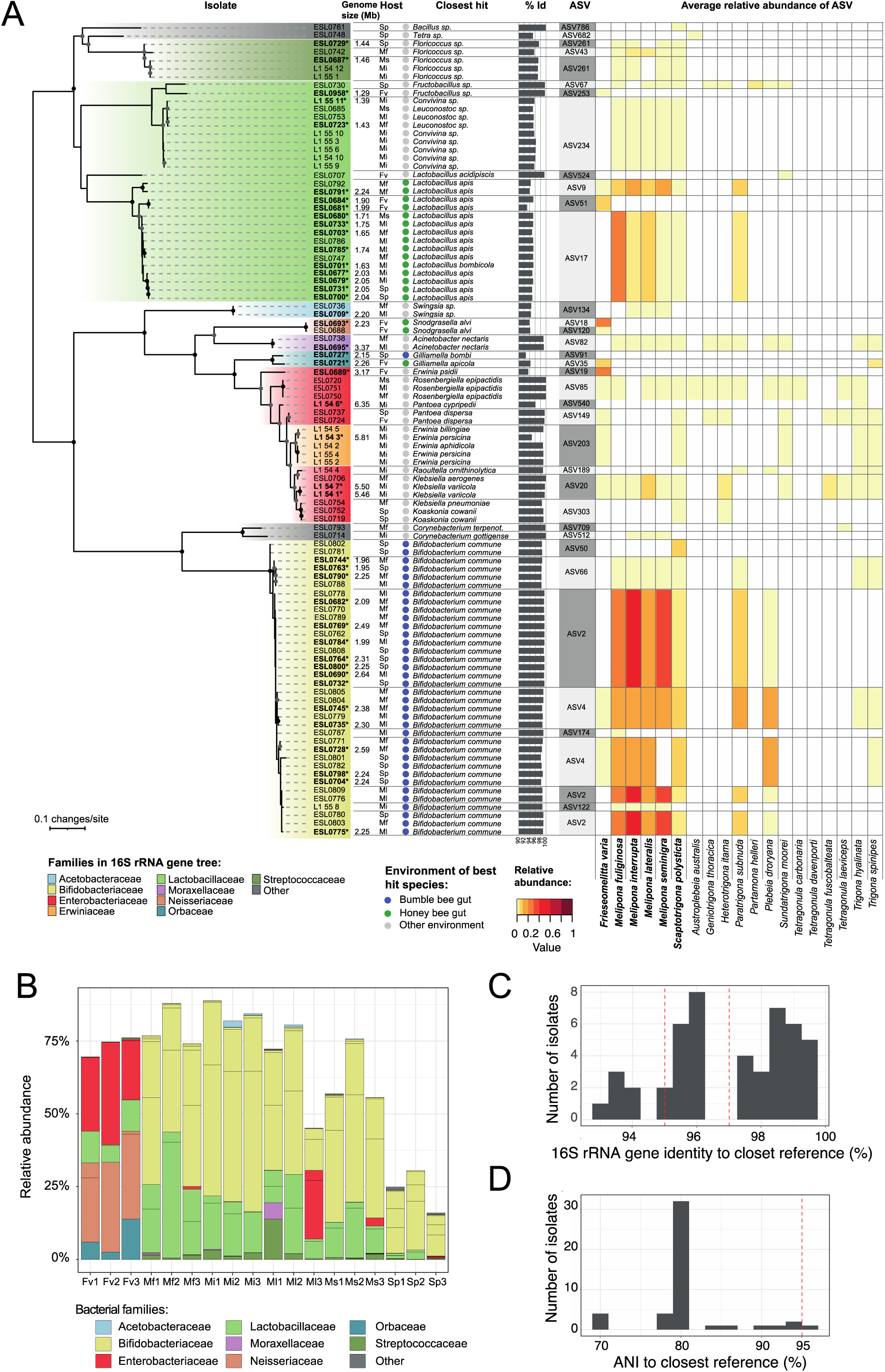
Bacterial strains isolated from the gut of six stingless bee species from Brazil. **(A)** Maximum likelihood tree was inferred from nearly complete sequences of the 16S rRNA gene of each isolate. Strain names of isolates for which we sequenced the genome are indicated in bold. The genome size of each sequenced strain is noted next to it. The isolate host is indicated as follows: Fv – *Frieseomelitta varia,* Ms – *Melipona seminigra*, Ml – *Melipona lateralis*, Mf – *Melipona fuliginosa*, Mi – *Melipona interrupta*, Sp – *Scaptotrigona polysticta*. Closest hit indicates best BlastN hit of the 16S rRNA gene sequence against the 16S ribosomal RNA (NCBI: Bacteria and Archaea type strains) database. Colored circles indicate if the strain of the best hit was isolated from the gut of a bumble bee, a honey bee, or from elsewhere. Bar plots indicate % identity of the best BlastN hit. The matching ASV and its average relative abundance across the nineteen analyzed stingless bee species are indicated. The names of the stingless bee species from our study are given in bold. Note that many isolates matched the same ASV. **(B)** Relative abundances of matching ASVs in each of the 18 samples of the six stingless bees species sampled in our study, i.e. representativeness of the isolates in the amplicon data. **(C)** Distribution of the isolated strains based on best 16S rRNA gene identity to closest reference species. These are the same values as shown in Figure 2A. The dashed lines indicate 95% and 97% identity corresponding to the genus and species threshold, respectively. **(D)** Distribution of the isolated strains based on pairwise ANI with the closest reference genome publicly available. The dashed line indicates the species threshold at 95% ANI.

### Stingless bee isolates form deep-branching phylogenetic lineages related to bacteria isolated from honey bees and bumble bees

To assess the phylogenetic placement of the isolated stingless bee gut bacteria relative to gut bacteria from honey bees and bumble bees, we selected 46 strains from 10 different bacterial families for genome sequencing (**Figure 2A, Supplementary Table 4**). Using a combination of Illumina and Oxford Nanopore sequencing, we obtained 23 complete and 23 draft genomes (2-66 contigs). Genome size of the cultured isolates ranged from 1.2-6.3 Mb. *Fructobacillus* ESL0730 (1.2Mb) and the two Streptococcaceae strains ESL0687 and ESL0729 (1.4 Mb) harbored the smallest, and Leuconostocaceae ESL0723 the largest (6.3 Mb) genomes of the sequenced strains (**Supplementary Table 4**). Genome comparisons with other bacteria, including strains isolated from honey bees and bumble bees, showed that most stingless bee gut bacteria had 80% average nucleotide identity with previously sequenced strains indicating that we have isolated strains of novel bacterial species or genera (**Figure 2D**). Accordingly, genome-wide phylogenies based on single-copy orthologs showed that most isolates formed deep-branching, stingless bee-specific lineages, exclusive of any previously sequenced strain. However, consistent with the results of the 16s rRNA gene analysis, several of these lineages were related to major phylotypes of the honey bees and bumble bee gut microbiota (**Figure 3A-D, Supplementary Figure 2–6**) such as *Snodgrassella*, *Gilliamella*, *Lactobacillus* Firm-5, *Bifidobacterium*, and *Bombella*. In the case of *Snodgrassella*, *Gilliamella*, and *Lactobacillus* Firm-5, the stingless bee-specific lineages formed a monophyletic clade with lineages of honey bee and bumble bee isolates (**Figure 3A-C** and **3F**). Notably, in all three cases, the bacteria from stingless bees presented the earliest branching lineages, i.e. the honey bee and bumble bee gut bacteria diverged after the split from the stingless bee gut bacteria. While these results suggest that these bacteria have derived from a common ancestor that was already adapted to social bees, the bacterial phylogenies were incongruent with current phylogenies of the host, which show that the honey bees (Apini) diverged before the split of stingless bees (Meliponini) and bumble bees (Bombini) (**Figure 3E**). A different pattern was observed for *Bifidobacterium*. In this case, strains isolated from stingless bees, honey bees, and bumble bees were not monophyletic. In fact, the stingless bee isolates belonged to a different clade than the honey bee isolates, while the strains isolated from bumble bees belonged to either of them (**Figure 3D** and **3G**). Similarly, the two Acetobacteraceae strains (ESL0695 and ESL0709) were not monophyletic with the honey bee isolates of the genus *Bombella*, although they belonged to the same Hymenoptera-associated clade within this family (**Supplementary Figure 3**). This suggests that in both cases, *Bifidobacterium* and Acetobacteriaceae, bacteria of distinct lineages have independently adapted to the gut environment of social bees.

**Figure 3.**
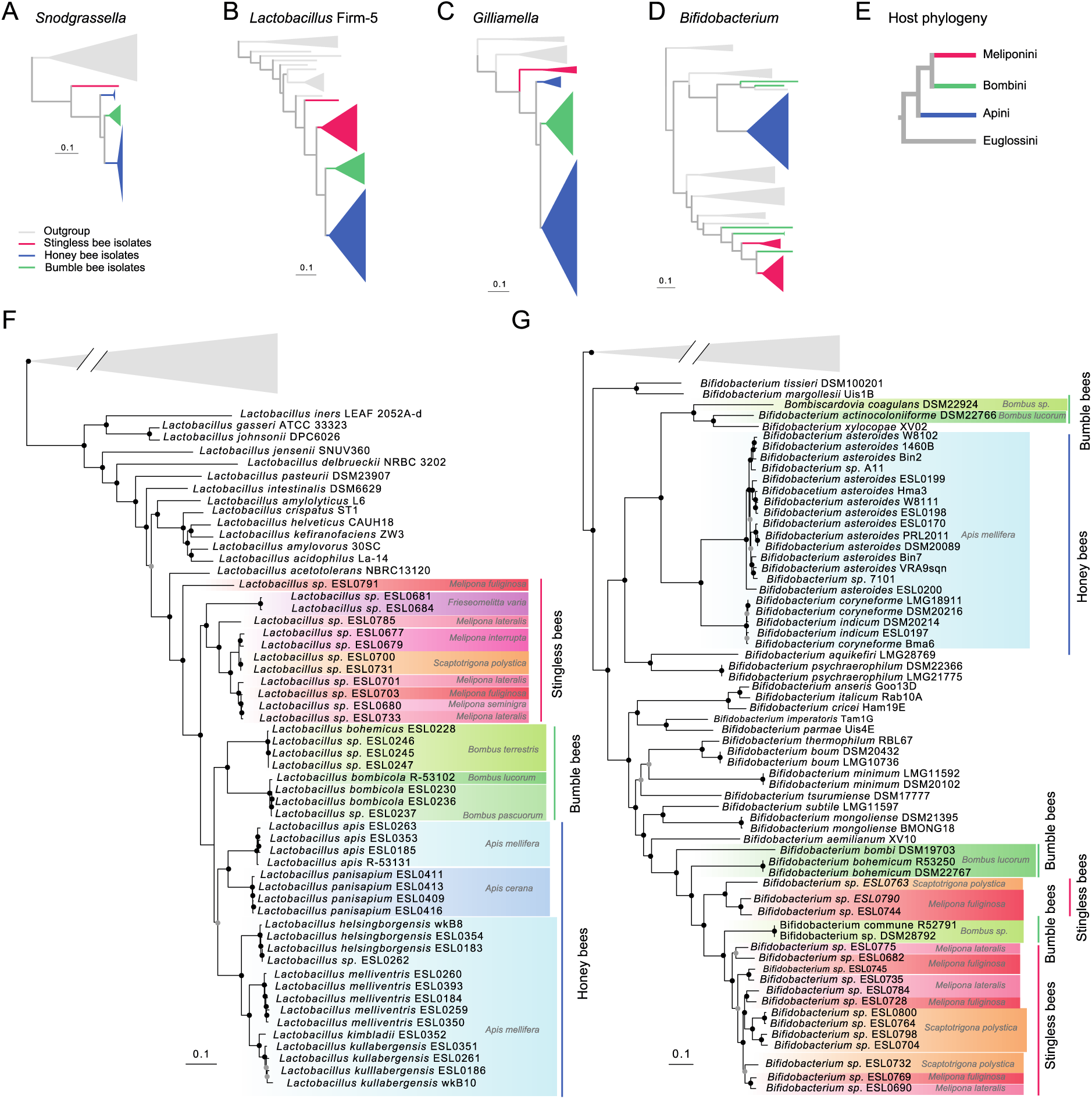
Isolates of stingless bee gut bacteria present novel species and belong to deep-branching phylogenetic lineages. **(A-D)** Simplified genome-wide maximum likelihood phylogenies of *Snodgrassella*, *Lactobacillus* Firm-5, *Gilliamella*, and *Bifidobacterium* based on single copy gene orthologs. All branches shown are supported by >95/100 bootstrap replicates. Most of the branches have been collapsed. **(E)** Dendrogram depicting the topology of the social bee phylogeny (adapted from (53, 54)). **(F)** Detailed genome-wide phylogeny of the genus *Lactobacillus* with bacteria belonging to the social bee-specific phylotype *Lactobacillus* Firm-5 highlighted in different colors according to the host species/group. The maximum likelihood tree was computed on the concatenated amino acid sequences of 355 single copy core genes using the substitution model LG+F+I+G4. **(G)** Detailed genome-wide phylogeny of the genus *Bifidobacterium* with bacteria belonging to social bee-specific clades highlighted in different colors according to the host species/group. The maximum likelihood tree was computed on the concatenated amino-acid sequences of 151 single copy core genes using the substitution model LG+F+I+G4.

Of the 46 strains selected for sequencing, 13 were not directly related to bacteria previously isolated from social bees. Most of these isolates matched to minor ASVs in our community profiling analysis, with the exception of three strains (ESL0689, ESL0687, and ESL0729, **Figure 2A**). ESL0689 corresponded to the Enterobacteriaceae ASV19 which dominated the communities of all three Fv samples (20-35% of the reads per sample). This isolate was situated on a long branch diverging between the genera *Klebsiella* and *Raoultella* (**Supplementary Figure 4**). ESL0687 and ESL0729 corresponded to ASVs of the family Streptococcaceae which were detected in several stingless bee species in our studies as well as in previous studies (20, 37, 38). They formed a deep-branching sister clade of the flower-associated genus *Floricoccus* (**Supplementary Figure 5A**). All three strains seem to be novel species based on their divergence to previously sequenced bacteria and may represent specialized gut symbionts of stingless bees.

### Stingless bee gut bacteria have diversified into distinct species and reveal a high extent of genomic diversity

Stingless bee isolates belonging to the same lineage were often separated by long branches in our phylogenies indicating substantial genomic divergence (**Figure 3F** and **G**, **Supplementary Figures 2–6**). This was confirmed by comparing pairwise 16S rRNA gene identity to genome-wide average nucleotide identity (ANI) between isolates of the same bacterial family. Despite high similarity in 16S rRNA gene identity, ANI was often <95% suggesting that most lineages of stingless bee gut bacteria contain several divergent species (**Figure 4A**). For example, the 12 sequenced strains of *Lactobacillus* Firm-5 fell into 8 distinct species-level clusters (i.e. ANI <95%, **Figure 4B**). A similar pattern was observed for the 17 *Bifidobacterium* strains, which fell into 14 distinct species-level clusters (**Figure 4C**), as well as the two *Gilliamella* and the two Streptococcaceae strains, which both also fell below the species-level ANI cut-off. Notably, some *Bifidobacterium* and *Lactobacillus* Firm-5 strains that were isolated from the same sample belonged to different ANI clusters, indicating that divergent bacterial species can co-occur in the same host species and colony. Inversely, strains belonging to the same ANI cluster were sometimes isolated from different bee species, suggesting that these bacterial species clusters are not necessarily host-specific (**Figure 4B** and **C**).

**Figure 4.**
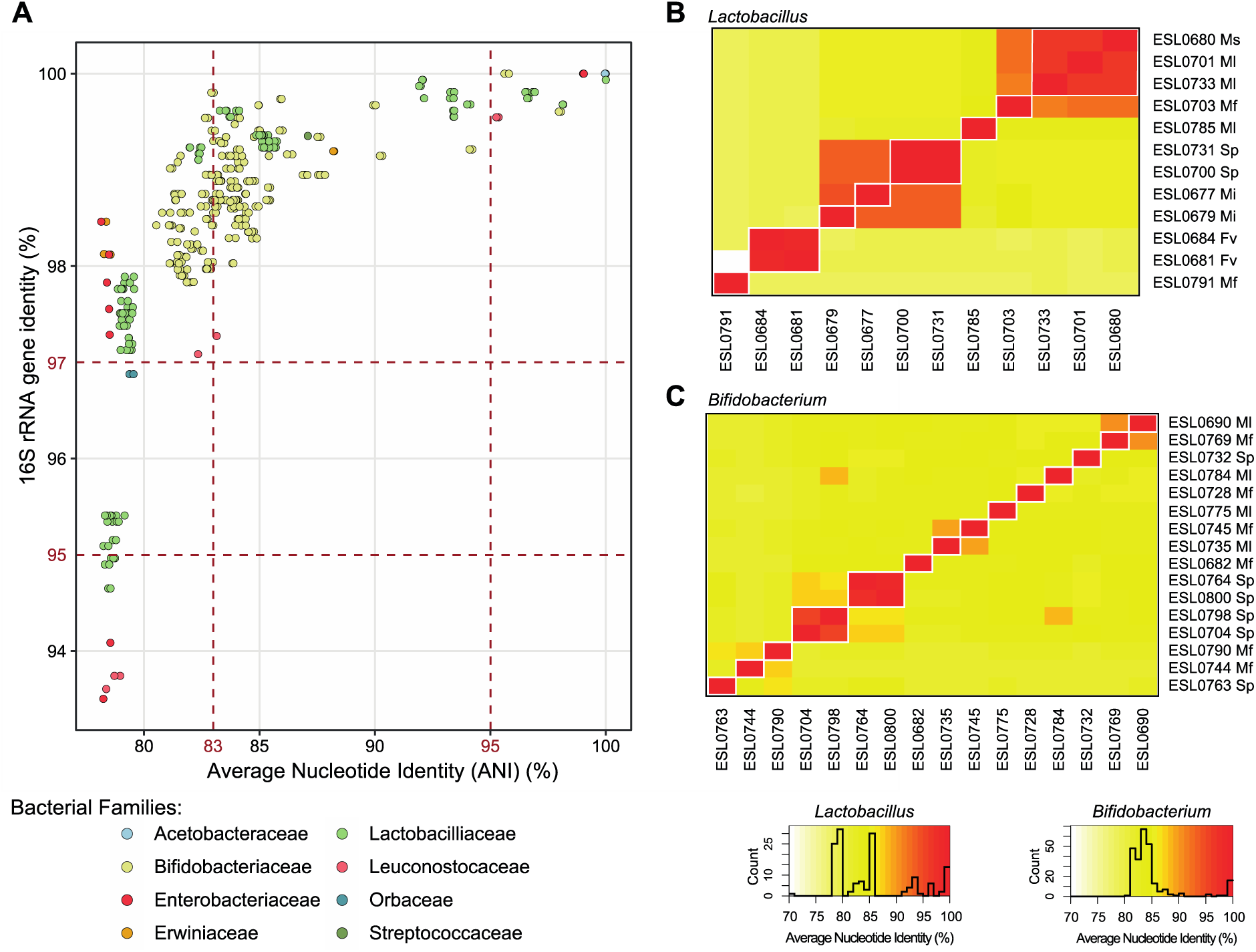
Genomic divergence among isolates with high 16S rRNA gene similarity. **(A)** The average nucleotide identity (ANI) versus 16S rRNA gene identity was plotted for each pair of isolate genomes belonging to the same bacterial family. The two vertical dashed bars indicate the ANI thresholds of 95% and 83% demarcating the intra-species and inter-species ANI values, respectively (65). The horizontal dashed lines delineate the commonly used species (>97%) and genus (>95%) thresholds for the 16S rRNA gene similarity. Average nucleotide identity heatmaps of **(B)** *Lactobacillus* Firm-5 isolates and **(C)** *Bifidobacterium* isolates. The 95% ANI clusters are delimited in white. The isolate host is indicated, Fv – *Frieseomelitta varia,* Ms – *Melipona seminigra*, Ml – *Melipona lateralis*, Mf – *Melipona fuliginosa*, Mi – *Melipona interrupta*, Sp – *Scaptotrigona polysticta*.

### Core microbiota members in stingless bees have similar functional capabilities as in honey bees and bumble bees

To assess the functional potential of stingless bee gut bacteria, we determined the genomic completeness of different metabolic pathways and functions in the genomes of the sequenced strains and compared it to related bacteria isolated from honey bees, bumble bees, or from elsewhere, and which were included in our phylogenomic analysis. We specifically looked at energy and carbon metabolism, amino acid, co-factor, and nucleoside biosynthesis, as well as secretion, motility, and adhesion (**Figure 5**).

**Figure 5.**
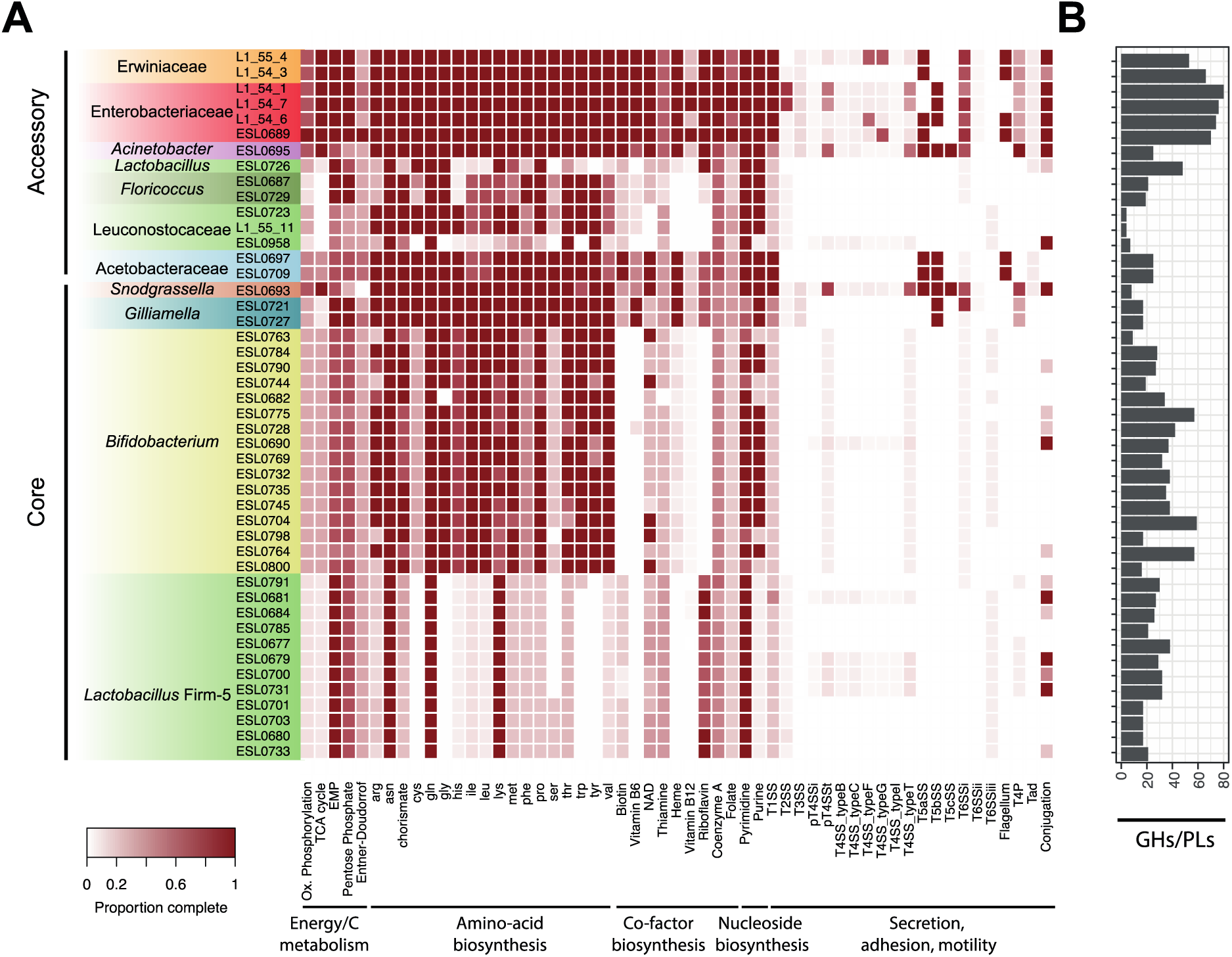
Metabolic capabilities of the sequenced stingless bee gut isolates. **(A)** The heatmap indicates the genomic completeness of major metabolic pathways and functions related to energy and carbon metabolism (EMP: Embden–Meyerhof–Parnas), the biosynthesis of amino-acid, co-factors, and nucleoside, as well as secretion, adhesion, and motility across the sequenced bacterial isolates. The isolates are ordered into core and accessory members based on whether they are related to bacteria isolated from other social bees. **(B)** Total number of glycoside hydrolase (GH) and polysaccharide lyase (PL) genes in each isolate.

#### Energy and carbon metabolism

Many of the sequenced strains (39/46) - including all Lactobacillaceae, the Bifidobacteria, and the two Streptococcaceae and *Gilliamella* strains – were missing key genes of the TCA cycle and for oxidative phosphorylation, but encoded functions for the breakdown (mostly glycoside hydrolases (GH)), and oxidation of sugars (Embden-Meyerhof-Parnas (EMP), pentose phosphate (PPP), and/or Entner-Doudorrof (ED) pathways). A detailed analyses of the enzymes used for carbohydrate breakdown by the core members *Lactobacillus* Firm-5, *Bifidobacterium*, and *Gilliamella* showed that the stingless bee gut bacteria encoded similar glycoside hydrolase family genes as honey bee and bumble bee isolates (**Figure 6**), including enzyme families for cleaving plant-derived glycans. For example, glycoside families GH5, GH30, GH31, GH42, GH43, and GH51 are involved in the degradation of hemicellulose (27, 30, 52). GH78 can be responsible for the cleavage of rhamnose residues from rutin, a major pollen-derived flavonoid that was demonstrated to be deglycosylated by honey bee isolates of *Lactobacillus* Firm-5 which encode GH78 genes (52). Another example is GH13 which includes neopullulanases and α-amylases for the breakdown of plant-derived starch. Together, these results suggest that stingless bee gut isolates of *Lactobacillus* Firm-5, *Bifidobacterium*, and *Gilliamella* are saccharolytic fermenters that breakdown pollen or nectar-derived glycans, similar as previously reported for the corresponding bacteria in the gut of honey bees or bumble bees. Notably, there was substantial variation in the number and type of glycoside hydrolase family genes between divergent strains, which goes in line with the extensive genomic diversity detected between stingless bee gut isolates of these three phylotypes. A complete TCA cycle was only found in the genomes of Neisseriaceae sp. ESL0693, *Acinetobacter* ESL0695, the Enterobacteriaceae, and the Erwiniaceae. The same strains also harbored the most complete gene sets for oxidative phosphorylation. Notably, Neisseriaceae sp. ESL0693 also lacked key genes in the EMP, PPP, and the ED pathways and contained very few GH family genes (**Figure 5**). This suggests that this bacterium cannot utilize sugars and obtains energy via aerobic respiration, as previously found for *Snodgrassella* isolates of honey bees and bumble bees (23).

**Figure 6.**
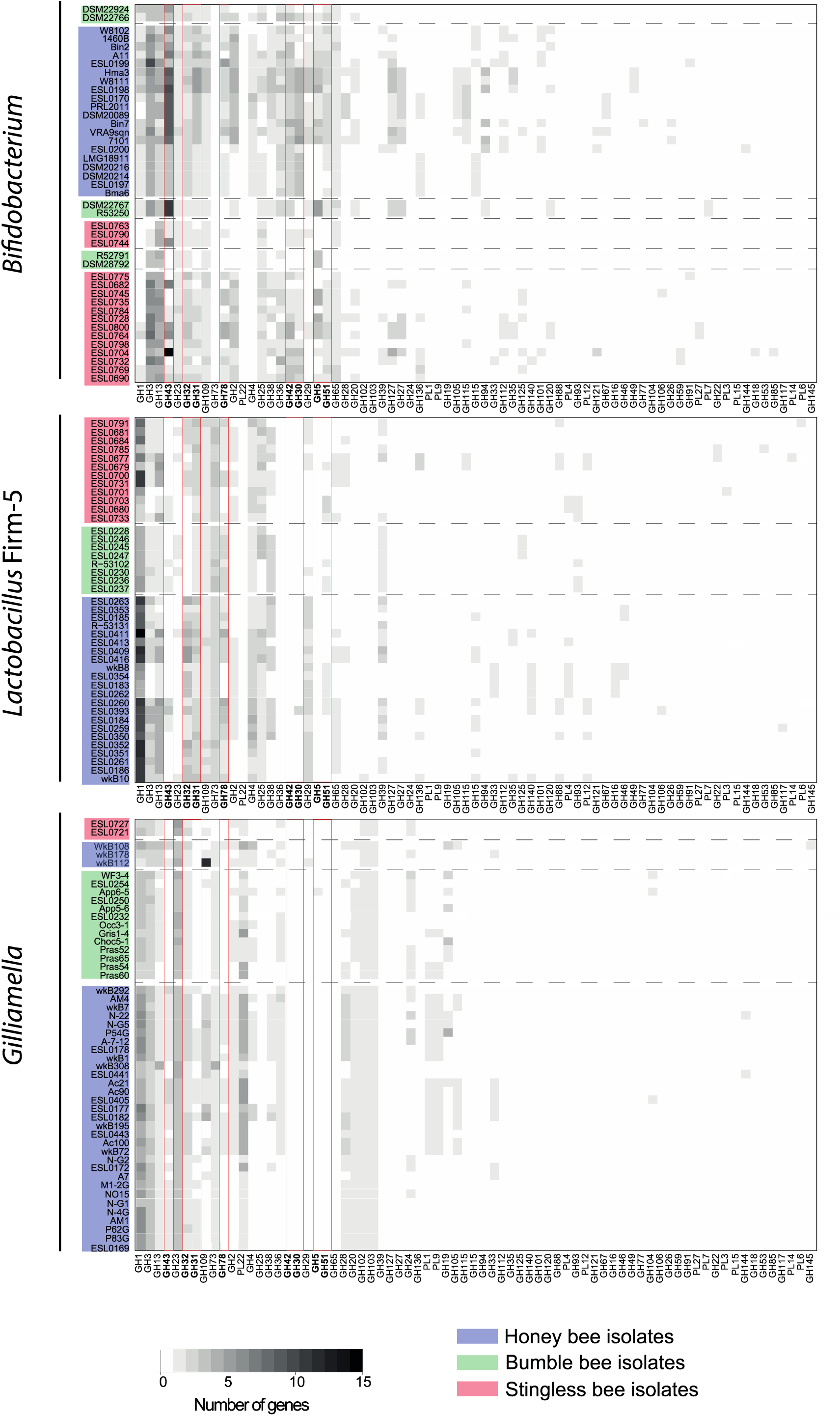
Carbohydrate-active enzyme profiles for *Bifidobacterium, Lactobacillus* Firm-5 and *Gilliamella* isolates. Distribution of genes in each GH and PL families respectively for *Bifidobacterium, Lactobacillus* Firm-5 and *Gilliamella* isolates. The isolates were sorted according to their position in the phylogenies, the hosts are indicated by the background colors. Glycoside hydrolases families mentioned in the text are outlined with a red square. The color scale indicates the number of genes in each family.

#### Amino acid, nucleoside, and co-factor biosynthesis

Differences between stingless bee gut isolates of different taxonomic groups were also found in terms of their biosynthetic potential. Strains belonging to the *Lactobacillus* Firm-5 clade were auxotrophic for the production of most amino acids (i.e. all except for Lys, Gln, and Asn) as well as purine and several co-factors (e.g. heme, vitamin B6 and B12) (**Figure 5**). Isolates of the Bifidobacteria, Streptococcaceae, and Leuconostocaceae were also auxotrophic for many co-factors, but for much fewer amino acids than *Lactobacillus* Firm-5. Interestingly, there was variation in auxotrophies among the Bifidobacterial strains, especially for the production of purine, NAD+, Thr, Lys, Arg, Gly, and chorismate. Other strains (such as those of *Gilliamella*, *Snodgrassella*, Acetobacteraceae, Enterobacteriaceae, and Erwiniaceae) had fewer auxotrophies. Similar biosynthetic capability profiles were found in related strains included in our phylogenies, which suggests that these functional profiles are not specific to stingless bee gut bacteria but rather conserved across the entire phylotype (**Supplementary Figures 2–6**).

#### Secretion, adhesion, motility

Secretion systems, pili, and flagella were mostly restricted to the gram-negative bacteria of the isolated strains. Type 1, Type 5, and Type 6 secretion systems were prevalent across these bacteria, whereas Type 2 and Type 4 secretion systems were only present in a few strains (**Figure 5**). Flagella were detected in the two Acetobacteraceae, all Erwiniaceae, and Enterobacteriaceae strain ESL0689. Tad pili were not detected in any of the analyzed bacteria, while Type IV pili components were mostly found in Neisseriaceae sp. ESL0693 and *Acinetobacter* ESL0695, and to some extent also in Orbaceae, Erwiniaceae, and Enterobacteriaceae ESL0689. Similar gene sets for secretion, adhesion, and motility were also found in related gut bacteria from honey bees or bumble bees, as shown by the functional profiles of all strains included in our phylogenies (**Supplementary Figures 2–6**).

Altogether, this first assessment of the gene content of the stingless bee gut bacteria show that they have similar functional potential as isolates from honey bees and bumble bees suggesting that they occupy similar ecological niches in the gut across social bees.

## Discussion

Previous findings suggested that the core members of the bee gut microbiota have been acquired in a common ancestor of the social bees (20) and possibly co-diversified with their hosts over millions of years (16, 33). However, the lack of genomic data from gut bacteria of stingless bees has limited our view on the evolution of these specialized microbial communities. With the establishment of isolate genomes of diverse stingless bee gut bacteria our study fills an important knowledge gap and provides new lines of evidence that rule out strict co-diversification between the core microbiota members and social bees.

Our genome-wide phylogenies of *Snodgrassella*, *Lactobacillus* Firm-5, and *Gilliamella* revealed that the identified lineages of stingless bee gut bacteria branched off before the divergence of the lineages of honey bee and bumble bee gut bacteria. This pattern is not congruent with the topology of the host phylogeny, in which honey bees diverged before the split of stingless bees and bumble bees (53, 54). The basal split of the stingless bee gut bacterial lineages depends on the rooting of our trees being correct. However, all phylogenies were supported by high bootstrap values at the critical nodes suggesting robust phylogenetic signals in our datasets. Our phylogenies also showed that for both *Gilliamella* and *Snodgrassella*, honey bee isolates were not monophyletic, i.e. some lineages branched before and others after the divergence of the bumble bee clades, which is inconsistent with co-diversification. This was already noted in a previous study (33), and similar results were also obtained for Lactobacillus Firm-5 based on the phylogenetic analysis of single protein-coding genes cloned from different social bee species (20). Finally, our phylogenomic analysis showed that honey bee and stingless bee isolates of *Bifidobacterium* belonged to two separate clades of the Bifidobacteriaceae suggesting that these bacteria have independently adapted to the gut environment of social bees.

Co-diversification can only occur when symbionts and hosts exhibit a high degree of partner fidelity and are transmitted vertically from one generation to the next over many generations (55). While most core microbiota members of social bees indeed seem to have a host-restricted distribution (20), examples of lineages with a broader host range exist as well (56). Moreover, some strains have been experimentally shown to be able to colonize non-native hosts demonstrating that host jumps are possible (20, 27). Our study showed that closely related stingless bee species (i.e. from the same or related genera) can have overlapping community profiles with predominant ASVs being shared across hosts. Similar observations have been made in other 16S rRNA gene profiling studies of stingless bees (20, 37). While such analyses often provide insufficient resolution to discriminate between closely related strains or species, our genomic analyses confirmed that stingless bees isolates of *Bifidobacterium* and *Lactobacillus* do not necessarily cluster by host species. It is possible that the strength of host specificity varies across social bees, or bacterial lineages, depending on the symbiotic function of the gut bacteria, or the hosts’ divergence, ecology, or geographic distribution. This would influence the extent to which gut bacteria can co-diversify within certain host lineages. To test this hypothesis, future studies would have to compare the strength of host specificity across multiple bees using representative sets of host species in each of the tree main bee clades.

Another piece of evidence indicating that the microbiota across social bees may be more variable than previously assumed comes from the observation that some of the designated core members were not always detected in the sampled bees. For example, while *Lactobacillus* and *Bifidobacterium* were prevalent across all six stingless bee species sampled in our study, they were absent from the gut microbial communities of some of the previously sampled bee species (15, 17, 37, 57). Likewise, stingless bees of the genus *Melipona* were shown to systematically lack the two core members *Snodgrassella* and *Gilliamella* (37). Both taxa were also rare across the four *Melipona* species analyzed in our study. However, three out of 12 analyzed colonies had high abundances of *Snodgrassella*. This suggests that this bacterium is not completely absent from this bee genus, but may be occasionally acquired from other bee species, varies in prevalence depending on season, bee age, or development, or is restricted to only the *Melipona* species analyzed in our study.

Finally, representatives of core members of the social bee gut microbiota were recently also found in bees of the distant genus *Xylocopa* (Carpenter bees) (58, 59), suggesting that these bacteria may have been associated with bees before the emergence of sociality, or that they have a broader and less-specific distribution across bees than previously suspected.

In summary, our results together with previous findings indicate a rather dynamic evolutionary background of the core members of the social bee gut microbiota. Rather than having strictly co-diversified with their hosts extended periods of host-restricted evolution (and likely co-diversification in some lineages) seem to have been interrupted by host switches, and independent symbiont gains and losses. Our observation that the stingless bee isolates repeatedly form the most a sister group to bumble bee and honey bee isolates (for *Gilliamella*, *Snodgrassella*, and *Lactobacillus* Firm-5) is intriguing given the host phylogeny. It may suggest that these core members have an origin in stingless bees and then spread to the other two groups, especially for *Lactobacillus* where two stingless bee isolates clades split before the split between bumble bee and honey bee isolates. However, given the large diversity of social and solitary bees, it is clear that the currently available datasets are insufficient to explain the distribution and phylogenetic relationships of these gut symbionts across hosts. Broader samplings of stingless bees, honey bees, and bumble bees, combined with genome-resolved approaches, are needed to fully understand the diversity, distribution, and evolutionary trajectories of social bee gut bacteria and to accurately reconstruct the ancestral bee microbiome composition. Formal analysis should be applied to test for co-diversification.

The importance of sampling biases when assessing patterns of co-diversification between hosts and their gut bacteria is highlighted by the analyses of Bacteroidaceae gut symbionts of hominids. While originally reported to have co-diversified with their hosts (60), re-examination with increased sampling disrupted the co-diversification pattern observed earlier (61). In contrast, a recent study identified strong signals of parallel evolutionary history between seven (out of 56 tested) gut bacterial taxa and human populations (62), and phylogenetic congruency has also been found for certain stinkbug insects and their primary gut symbiont (63). This demonstrates that co-diversification has occurred between certain gut bacteria and their hosts.

Besides offering new insights into the evolution of the social bee gut microbiota, our genomic analysis also revealed the functional potential of major gut symbionts of the analyzed stingless bee species. All isolates of *Lactobacillus* Firm-5, *Bifidobacterium*, and *Gilliamella* carried genes for the saccharolytic fermentation of diet-derived carbohydrates. In contrast, *Snodgrassella* ESL0689 lacked such functions in its genome, but instead harbored genes for aerobic respiration. These results are consistent with findings from honey bees and bumble bees (22, 23, 26–28, 30) and hence suggest that the core microbiota members occupy similar ecological niches across the three groups of social bees.

Another parallel to findings from honey bees and bumble bees was the extensive genomic divergence present among strains of the core members *Lactobacillus* Firm-5, *Gilliamella*, and *Bifidobacterium*, even when isolated from the same host species. Moreover, we found genomic variation in carbohydrate breakdown and amino acid and nucleoside biosynthesis functions among these strains. This suggests that the diversification of these bacteria has not only been driven by isolation into different host species but also by the adaptation to different ecological niches in the gut, similar as shown for bumble bees and honey bees (28, 33, 35, 64). These parallels may not be surprising as the dietary preferences of the analyzed stingless bee species are similar to those of honey bees and bumble bees. In the future, it will be interesting to look at the functional potential of the core microbiota members in bees with different dietary habits, such as the vulture bees (38), i.e. stingless bees that feed on raw meat instead of pollen yet share a subset of the core members with other social bees.

Some of the isolate genomes we sequenced in our study did not come from any of the core members of the social bee gut microbiota. They may present transient community members, opportunistic pathogens, or host-specific gut symbionts with complementary functions to the core microbiota. Therefore, the established genomes present an important resource for future research. A particular strain that drew our attention was Enterobacteriaceae ESL0689, as it belonged to an ASV that was present at high relative abundance in all three colonies of *Fv.* ESL0689 harbored a complete TCA cycle and a respiratory chain and could also synthesize most amino-acids and co-factors indicating a similar metabolic niche as *Snodgrassella* (23).

In conclusion, our study provides new insights into the evolution of the social bee gut microbiota and presents a first step in characterizing the functional potential of major gut bacteria present in stingless bees. However, given the large diversity of stingless bees with hundreds of different species distributed throughout the tropical and subtropical regions of the world, it is clear that our study only presents the starting point in characterizing their genomic diversity and functional potential. More detailed studies and larger genomic survey combined with experimental analyses will be needed to understand their evolution and assess their impact on the host.

## Methods

### Bee sampling

Bees were collected from three different nests of each of the following six Meliponini species in February and March 2019: *Frieseomelitta varia*, *Scaptotrigona polysticta*, *Melipona fuliginosa*, *Melipona interrupta*, *Melipona lateralis* and *Melipona seminigra*. All nests were located in a rural meliponary in the vicinities of Iranduba municipality (Iranduba-AM, Brazil, 3°10’52.7”S 60°07’08.5”W), and the sampling was carried out under the SISGEN collection permission A256E82. Bees were sampled at the entrance of each nest and immediately immobilized by cooling at 4 °C. Then the entire gastrointestinal tract was dissected, and the hindgut separated from the anterior gut parts. Hindguts of bees from the same nest were pooled in two separate tubes. One pool per nest was mixed with 1x PBS and glycerol, homogenized using bead-beating, and subsequently cryopreserved at −80°C. This pool was used for bacterial culturing as described below. The other pool was cryopreserved at −80°C without homogenization and was subsequently used for DNA extraction and 16S rRNA gene analysis. For the sample used for sequencing, 20 bees per nest were pooled, while for the sample for culturing, only 3 bees per nest were pooled. For *Frieseomelitta varia* and *Scaptotrigona polysticta*, we pooled 40 and 10 bees, respectively, due to the small size of these bee species.

### Bacterial culturing

For establishing a culture collection of primary isolates from the gut of the six sampled bee species, serial dilutions of the cryopreserved homogenates were plated on eight different media: CBA (Columbia Blood Agar supplemented with 5 % defibrinated sheep blood (Thermofisher)), MRSA (De Man, Rogosa and Sharpe agar) supplemented with Fructose (2%) and L-Cysteine (0.1%), MRSA supplemented with Mannitol (2%), Chocolate Agar, TSA (tryptone soya agar), TYG (tryptone glucose yeast extract Agar), GC, LBA (Luria Bertani agar) without NaCl, BHIA (brain heart infusion agar) and SDA (Sabouraud dextrose agar). Plates were incubated in two different conditions: in a microaerobic 5% CO_2_-enriched atmosphere and in an anaerobic chamber (72% N_2_, 8% H_2_, 20% CO_2_), both at 34°C. After 2-7 days of incubation, colonies of different size and appearance were picked and re-grown on the same media and culturing conditions. Cryo-stocks of bacterial strains of interest were prepared by harvesting bacterial biomass in liquid media corresponding to the solid growth media and supplemented with 20% glycerol. For DNA isolation, bacteria were grown from the stocks, and a single colony was picked and re-grown on fresh media before harvesting bacterial biomass.

### Genotyping

All colonies that were selected for culturing were genotyped by PCR and Sanger sequencing of a 16S rRNA gene fragment. To this end, a small amount of bacterial material was transferred to a lysis buffer (Tris-HCl 1M pH=7.5, EDTA 0.5M, SDS 10%) containing 2.5 μl lysozyme (20mg/mL) and 2.5 μl proteinase K (20mg/mL), and incubated for 10 min at 37°C, 20 min at 55°C, and 10 min at 95°C. PCR was performed with universal bacterial primers that amplify the V1-V5 region of the 16S rRNA gene (27F - AGRGTTYGATYMTGGCTCAG and 907R-CCGTCAATTCMTTTRAGTTT) using the following reagents and thermocycler program: initial denaturing at 94°C for 5 min, followed by 32 cycles of denaturing at 94°C for 30 sec, annealing at 56°C for 30 sec, and extension 72°C for 1 min, and a final extension at 72°C for 7 min. PCR results were checked on a 1% agarose gel. PCR reactions selected for Sanger sequencing were purified using ExoSAP-IT^TM^ (1μl ExoSAP 5x, 4μl ddH_2_O) with the thermocycler program: 30 min at 37°C followed by 15 min at 80°C. Purified samples were then sent to Eurofins® for sequencing. Sanger sequences were analyzed with Geneious suite (Geneious®) and compared to GenBank at NCBI using BLAST tools (66).

### DNA isolation and genome sequencing

DNA isolation for Illumina sequencing was carried out using a customized SPRI bead-based extraction method or the FastPure Bacteria DNA Isolation Mini Kit (Vazyme). For the SPRI-bead method, bacteria were harvested and resuspended in tubes containing 200 mg of 0.1 mm-sized acid washed glass beads and 200 µl of TER buffer (10 mM Tris-HCl, 1 mM EDTA, 100 µg /ml RNAse A, pH8.0). Samples were homogenized using a FastPrep-25 5G instrument (2 rounds of 30 sec with the power set to 6) and subsequently centrifuged at max speed for 10 min at room temperature. 40 µl of SPRI beads were added to 100 µl of supernatant, immediately mixed thoroughly by repeated pipetting (>20x) and incubated for 5 min at room temperature. After placing the tubes on a magnet stand the liquid was removed and discarded and the beads washed two times with 200 µl 80% ethanol. After air drying the tubes on the magnetic stand 22 µl of 5 mM Tris-HCl pH8 was added. For isolating bacteria with the FastPure Bacteria DNA Isolation Mini Kit the manufacturer’s protocol for gram-positive bacteria was followed.

DNA isolation for Oxford Nanopore Technologies (ONT) sequencing was carried out using a custom DNA extraction protocol for Gram-positive bacteria. Tubes were prepared with glass beads and 160 μl of buffer P1 (Qiagen). Then bacteria were harvested and resuspended in these tubes by intensive vortexing. Lysozyme (20 μl, 100 mg/ml) was added and after gentle mixing, tubes were incubated at 56°C with shaking at 600 rpm for 30 min. Then 4 μl RNase A (100 mg/ml) were added to the tubes followed by 150 μl of buffer AL (lysis buffer, Qiagen). After mixing by vortexing, tubes were incubated in a thermomixer (37°C, 900 rpm) for 20 min. Tubes were centrifuged 10 min at 14’000 rpm to pellet the beads, the supernatant was transferred to new tubes with 35 μl Na-acetate and 270 μl isopropanol and mixed by inverting. Following an incubation of 1 h at 4°C, DNA was pelleted by centrifugation at 14’000 rpm for 10 min at 25°C. The supernatant was discarded, and the pellets were washed with 1 ml EtOH 80%. After a second centrifugation (14’000 rpm, 10 min at 25°C) the ethanol was removed, and the pellet left to dry at room temperature. DNA pellets were solubilized with 50 μl TER (10 mM Tris-HCl, 1 M EDTA, pH 8.0, 2 mg/ml RNase A) and tubes incubated at 37°C for 15 min. The solution was then transferred to PCR tubes. 40 μl NGClean beads were added, and the solution mixed by repeated pipetting. After a 5 min incubation, PCR tubes were placed on magnetic stands. When the solutions were clear, the liquid was removed, and the beads washed two times with 200 μl of 80% EtOH. Upon complete drying, beads were resuspended with 22 μl of 5mM Tris-HCl (pH 8.0). Finally, tubes were placed again on the magnetic stand and when the solution were clear, the supernatant was transferred to new 1.5 ml Eppendorf tubes.

Illumina sequencing libraries were prepared using the Nextera DNA Flex Library Preparation kit following instructions of the Illumina Reference Guide. This was followed by indexing, dilution, and denaturation according to the ‘Index Adapters Pooling Guide’ and the ‘MiniSeq System Denature and dilute Libraries guide’. Libraries were checked and quantified using a dsDNA fluorescent dye, before loading them on a MiniSeq High-output flowcell (150PE). ONT libraries were prepared using the ligation-based approach (LSK109). The sequencing was conducted on a ONT MinION for a duration of 72 hours with the high-accuracy base-calling using Guppy (*v5.0.11*).

### Genome assembly

Forty-six isolates from the stingless bees were sequenced with Illumina MiniSeq (150PE). Raw reads were checked using FastQC *v0.11.9* (67) and trimmed by Trimmomatic v0.39 (68) using parameters: PE-phred33 AllIllumina-PEadapters.fa:3:25:7 LEADING:9 TRAILING:9 SLIDINGWINDOW:4:15 MINLEN:60. For *de novo* assembly we used SPAdes (–careful option, *v3.15.2*) (69). For assembly quality control, reads were mapped back against the assembly with BWA *v0.7.17* (*70*), samtools *v1.12* (71) and plotted with R *v4.1.1*. Genomes completeness was evaluated with checkM *v1.0.13* (72).

Additionally, 26 isolates were also sequenced with Oxford Nanopore to produce long reads. Nanopore long reads were filtered with Filtlong *v0.2.0* (https://github.com/rrwick/Filtlong) for a minimum length of 7’000 and minimum mean q-score of 10 (min_length 7000, min_mean_q 10, length_weight 10). ONT-based assemblies were computed with Flye *v2.7.1* (73) over 5 iterations. Graphmap *v0.5.2* (74) and Racon *v1.0.1* (75) were used to perform two rounds of polishing. Finally, the racon-corrected assembly and Illumina reads were fed to Pilon *v1.24* (76) for single base and indels corrections.

### Genome annotation and analysis

Genomes were annotated with Prokka *v 1.13* (77). Phylogenies were computed for the bacterial families for which we had an isolate. For each family, we identified a set of closely strains and outgroup taxa and retrieved their genomes from NCIB and IMG/Mer (78). All genomes were re-annotated with Prokka to ensure annotations consistency. Gene orthology was inferred with OrthoFinder *v2.3.8* (*79*). Single copy ortholog genes were selected and their amino acid sequences were aligned (mafft *v7.453)* (*80*). An in-house script was used to trim the alignments by removing positions with more than 50% gaps and sequences belonging to the same genome were concatenated to produce a core gene alignment. This alignment was used to infer the maximum likelihood phylogeny using IQTree (*v1.7.beta17*, -st AA -bb 1000 -seed 12345 -m TEST) (81). For each phylogeny, the best evolutionary model was chosen according to BIC: *LG+I+G4* – Enterobacteriaceae; *LG+F+I+G4 –* Acetobacteraceae, *Bifidobacterium*, *Lactobacillus*, Leuconostocaceae, Moraxellaceae, Neisseriaceae and Streptococcaceae; and *JTTDMut+F+I+G4* – Orbaceae. Branch support of the trees were inferred using 1000 ultrafast bootstrap (UFBoot) repetitions. Clades can be trusted when UFBoot values are >95%.

### 16S rRNA gene based community profiling

Region V4 of the 16S rRNA gene was amplified using the primers 515F-Nex and 806R-Nex (TCGTCGGCAGCGTCAGATGTGTATAAGAGACAGGTGCCAGCMGCCGCGGTAA, GTCTCGTGGGCTCGGAGATGTGTATAAGAGACAGGGACTACHVGGGTWTCTAAT). The primers include Nextera XT index adapter sequences and the primers for the 16S rRNA V4 region (82) described by Kešnerová et al. (2020). The two step PCR was performed as following: the first PCR was performed using 12.5 µL of 2× Phanta Max Master Mix (Vazyme, Nanjing, China), 5 μl of MilliQ water, 2.5 μl of each primer (5 μM), and 2.5 μl of template DNA for a total volume of 25 μl. The PCR program started with a denaturation step at 98°C for 30sec, followed by 25 cycles of amplification (10 sec at 98°C, 20 sec at 55°C, and 20 sec at 72°C) and a 5 min final extension step at 72°C. The PCR products were verified by 2% agarose gel electrophoresis, then purified with clean NGS purification-beads in a 1:0.8 ratio of PCR product to beads and eluted in 27 μl Tris (10mM, pH 8.5). A second PCR step was performed to append the unique dual indexes to each sample in a total volume of 25 μl using 12.5 µL of 2× Phanta Max Master Mix (Vazyme, Nanjing, China), 5 μl of MilliQ water, 2.5 μl of Nextera XT index primers 1 and 2 (Nextera XT Index kit, Illumina) and 2.5 μl of templated DNA. The PCR program started with a denaturation step at 95°C for 3 min, followed by 8 cycles of amplification (30 sec at 95°C, 30 sec at 55°C, and 30 sec at 72°C), and a 5 min final extension step at 72°C. The libraries were then cleaned usingClean NGS purification beads (1:1.1 ratio of PCR product to beads) and were eluted in 27.5 μl Tris (10mM, pH 8.5). Prior to sequencing, the PCR product concentrations were quantified by PicoGreen and pooled in equimolar concentrations; the negative controls and blank extractions were pooled in equal volume. Sequencing was performed on an Illumina MiSeq sequencer (2 x 250bp) by the Genomic Technology Facility of the University of Lausanne. We followed DADA2 (83) pipeline to analyze the sequencing data. For the first part of the analysis, we executed the pipeline only on the 18 samples from our study. To control for possible contaminants, we used blank extractions and water as negative controls during the PCR. For the second part of the analysis, where we combined our dataset with the one from (20), we executed the pipeline independently a second time. Sequence quality control was performed with DADA2 integrated function *plotQualityProfile*. Data from the three sequencing runs were processed independently for the filtering, dereplication, the error rates calculation and the sample inference. After merging the denoised forward and reverse pairs, we merged the three sequence tables (*mergeSequenceTables*) and we applied the *collapseNoMismatch* function to unite similar ASVs with shifts or length variation. We selected sequences in range 250-256bp and removed chimeric sequences. Silva non-redundant SSU database v138.1 (83) was used for the taxonomic assignment of the ASVs. We filtered out 62 ASVs for which the taxonomic assignment was matching ‘Eukaryota’, ‘Chloroplast’ or ‘Mitochondria’. Finally, we removed samples with less than 5,000 reads. Non-metric MultiDimensional Scaling plots were computed in R using Bray-Curtis distances (Phyloseq: *ordinate*). The adonis package was used to compute permanovas.

### Functional Profiler

We developed a genomic profiler using several software to annotate the genomes and compute the completeness of key pathways and functions. The genomes were re-annotated with GhostKOALA (84) to get KEGG annotations of genes. We created a set of rules to define the steps of selected energy and metabolism pathways, and co-factors/nucleoside biosynthesis pathways and computed their completeness. In brief, an in-house script parsed the KEGG annotations of the genomes and for each pathways evaluated the completeness of each step. The pathways completeness was then summarized by counting the number of steps present relative to total number of steps needed. To obtain the completeness of amino acid biosynthesis pathways, we used GapMind (85). Secretion systems and related appendages were detected in the genomes by MacSyFinder’s TXSScan module (86). Finally, we ran dbCan (87) for the annotation of carbohydrate-active enzymes.

## Acknowledgements

We would like to thank Julien Marquis and the team from the Lausanne Genomics Technology Facility and Alban Ramette and his team from the University of Berne for carrying out the Illumina sequencing and Oxford Nanopore sequencing, respectively. We also would like to thank the ‘Ecole de Biologie’ of the University of Lausanne for their financial support to sequence the presented genomes in the context of the course “Sequence-a-genome” (SAGE) as part of the Master of Science in Molecular Life Science. Finally, we would like to thank all SAGE students of the years 2019-2020 and 2020-2021 for their contribution to this course: Yami Ommar Arizmendi Cárdenas, Samuel Aubert, Alec Auston, Fabrice Battiston, Etienne Bellani, Valentina Benigno, Valentin Borgeat, Alessandro Brandulas Cammarata, Marion Brechet, Marine Bugnon, Jessica Burnier, Théo Cavinato, Giacomo Ceracchini, Jérémy Cherbuin, Lucas Culebras, Audrey Daina, Hugues de Villiers de la Noue, Joe Dickinson, Alexandre Dudt, Elise Eray, Sara Ezzat, Christopher Forbes-Jaeger, Auriane Form, Léo Franchi, Pablo Guridi Fernández, Manon Henna, Aya Iizuka, Nicolas Jacquemin, Richie Kalusivikako, Maroussia Liechti, Simon Maréchal, Achille Mariotti, Aoife Mc Nally, Mam Malick Sy Ndiaye, Jade Nicolet, Astrid Oliva, Claire Paltenghi, Priyanka Parmar, Nicolas Pellaton, Katharina Pfaller, Brenda Ríos-Ochoa, Eric Risse, Daniel Rodriguez, Artemiy Saukin, Matthieu Simeoni, Miloš Stojanov, Marina Sudário, François Sutter, James Tan, Emanuele Tettamanti, Jamille Viray, Grazia Vizzarro and Chaymae Ziyani. This work was funded by the ERC Starting Grant (MicroBeeOme), the NCCR Microbiomes, a National Centre of Competence in Research, funded by the Swiss National Science Foundation (grant no. 180575) and a Swiss National Science Foundation Spirit grant (grant no. IZSTZ0_189496) to P.E.. F.M. was funded by the Swiss National Science Foundation (grant nr 315230_184908: SOMETALP).

## Data and materials availability

The bacterial genomes and sequencing data are available in the NCBI’s BioProjects <u>PRJNA906295</u> and <u>PRJNA897721</u>. Scripts used for the amplicon sequencing analysis, the Phylogenies and the metabolic profiler can be found on GitHub https://github.com/gsartonl/Publication_Sarton-Loheac_2022.

## Supplementary Figures legends

**Supplementary Figure 1.**
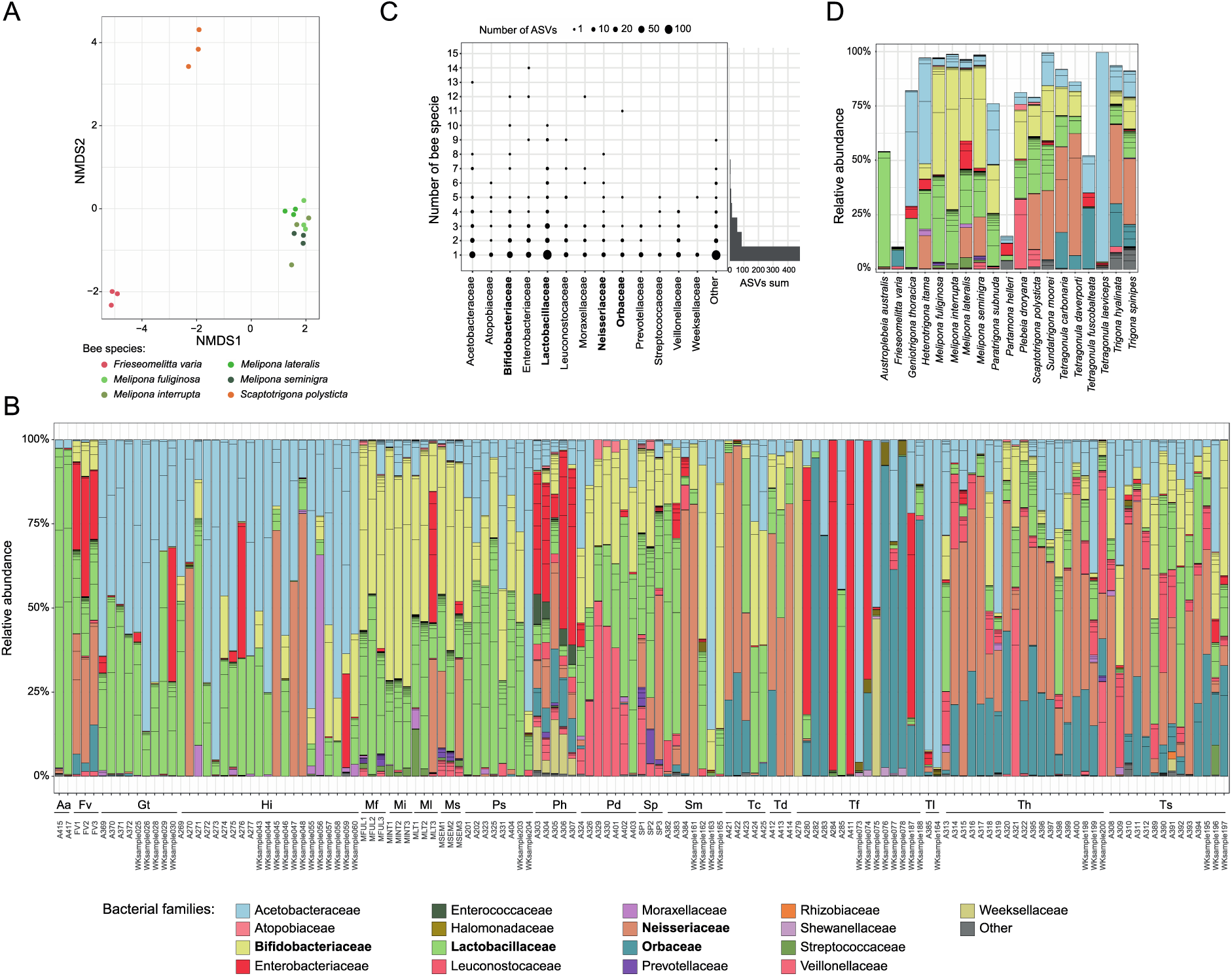
**(A)** NMDS based on ASV relative abundance in the 18 samples collected from six different stingless bee species in Brazil. The samples clustered by bee genus (PERMANOVA: ‘Genus’ pseudo F=11.05, p-value< 0.001). **(B)** Community profiles of the gut microbiota of 136 samples from 19 different stingless bee species collected in Brazil, Malaysia, and Australia. Relative abundance of ASVs is shown. ASVs are ordered and colored at the genus level according to the legend. ASVs with <1% relative abundances are summed up as ‘others’ and shown in grey. This plot includes the datasets from stingless bees from (20) and our study. The samples are ordered by bee species : Aa – *Austroplebeia australis*, Fv – *Frieseomelitta varia*, Gt – *Geniotrigona thoracica*, Hi – *Heterotrigona itama*, Mf – *Melipona fuliginosa*, Mi – *Melipona interrupta*, Ml – *Melipona lateralis*, Ms – *Melipona seminigra*, Ps – *Paratrigona subnuda*, Ph – *Partamona helleri*, Pd – *Plebeia droryana*, Sp – *Scaptotrigona polysticta*, Tc – *Tetragonula carbonaria*, Td – *Tetragonula davenporti*, Tf – *Tetragonula fuscobalteata*, Tl – *Tetragonula laeviceps*, Th – *Trigona hyalinata*, Ts – *Trigona spinipes*. **(C)** Number of shared ASVs. For each family, we reported how many ASVs were observed (indicated by the dot size) in how many bee species. Families with an average abundance <1% are grouped as ‘Other’. The sum of species-specific or shared (by two or more bee species) ASVs is reported on the right side. Core members families are highlighted in bold. **(D)** Relative abundances of the shared ASVs across the 19 stingless bee species (samples from the same species were pooled).

**Supplementary Figure 2.**
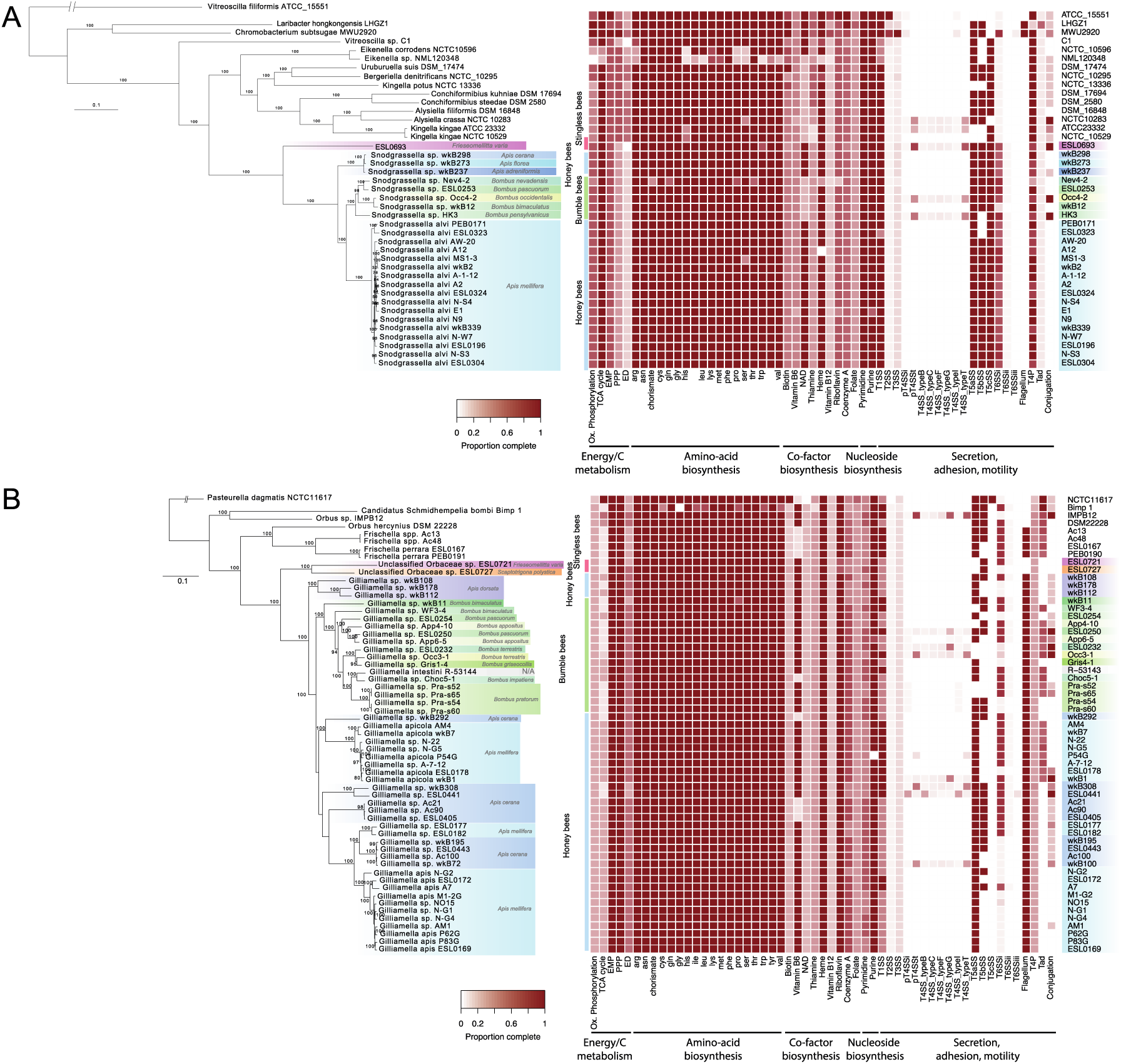
**(A)** Core genome phylogeny of the Neisseriaceae including phylotype *Snodgrassella*. The maximum likelihood tree was computed with the amino-acid sequences of 428 concatenated single copy core genes using LG+F+I+G4 substitution model. Bee isolates form a monophyletic clade. The heatmap indicates the genomic completeness of major metabolic pathways and functions related to energy and carbon metabolism (EMP: Embden–Meyerhof–Parnas), the biosynthesis of amino-acid, co-factors, and nucleoside, as well as secretion, adhesion, and motility across the sequenced bacterial isolates. **(B)** Core genome phylogeny of the Orbaceae including *Gilliamella* phylotype. The maximum likelihood tree was computed on the concatenated amino-acid sequences of 782 single copy core genes using JTTDCMut+F+I+G4 substitution model. The heatmap indicates the genomic completeness of major metabolic pathways and functions related to energy and carbon metabolism (EMP: Embden–Meyerhof–Parnas), the biosynthesis of amino-acid, co-factors, and nucleoside, as well as secretion, adhesion, and motility across the sequenced bacterial isolates.

**Supplementary Figure 3.**
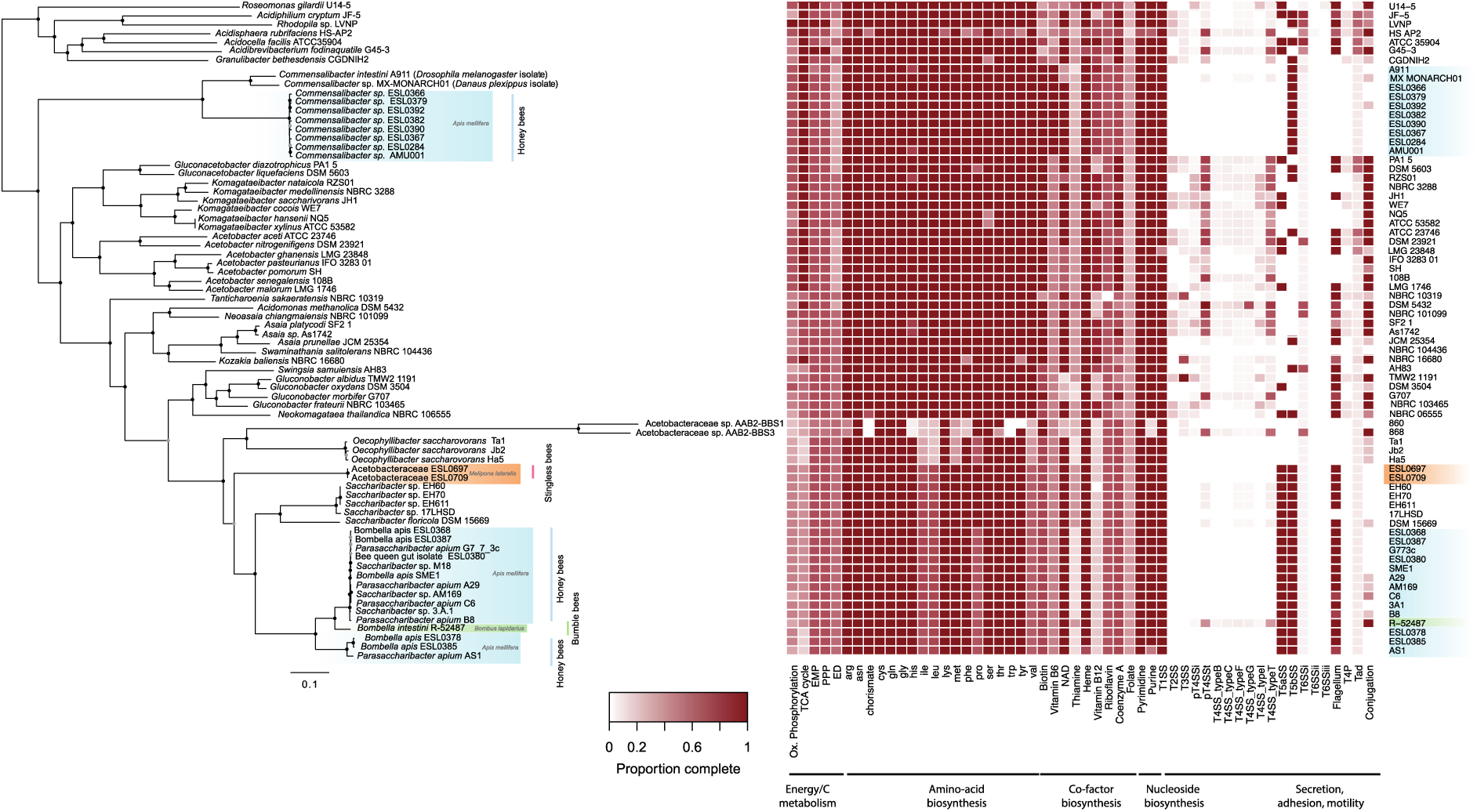
Core genome phylogeny of the Acetobacteraceae. The maximum likelihood tree was computed on the concatenated amino-acid sequences of 303 single copy core genes using LG+F+I+G4 substitution model. Bee isolates are distributed in two distant monophyletic clades. The heatmap indicates the genomic completeness of major metabolic pathways and functions related to energy and carbon metabolism (EMP: Embden–Meyerhof– Parnas), the biosynthesis of amino-acid, co-factors, and nucleoside, as well as secretion, adhesion, and motility across the sequenced bacterial isolates.

**Supplementary Figure 4.**
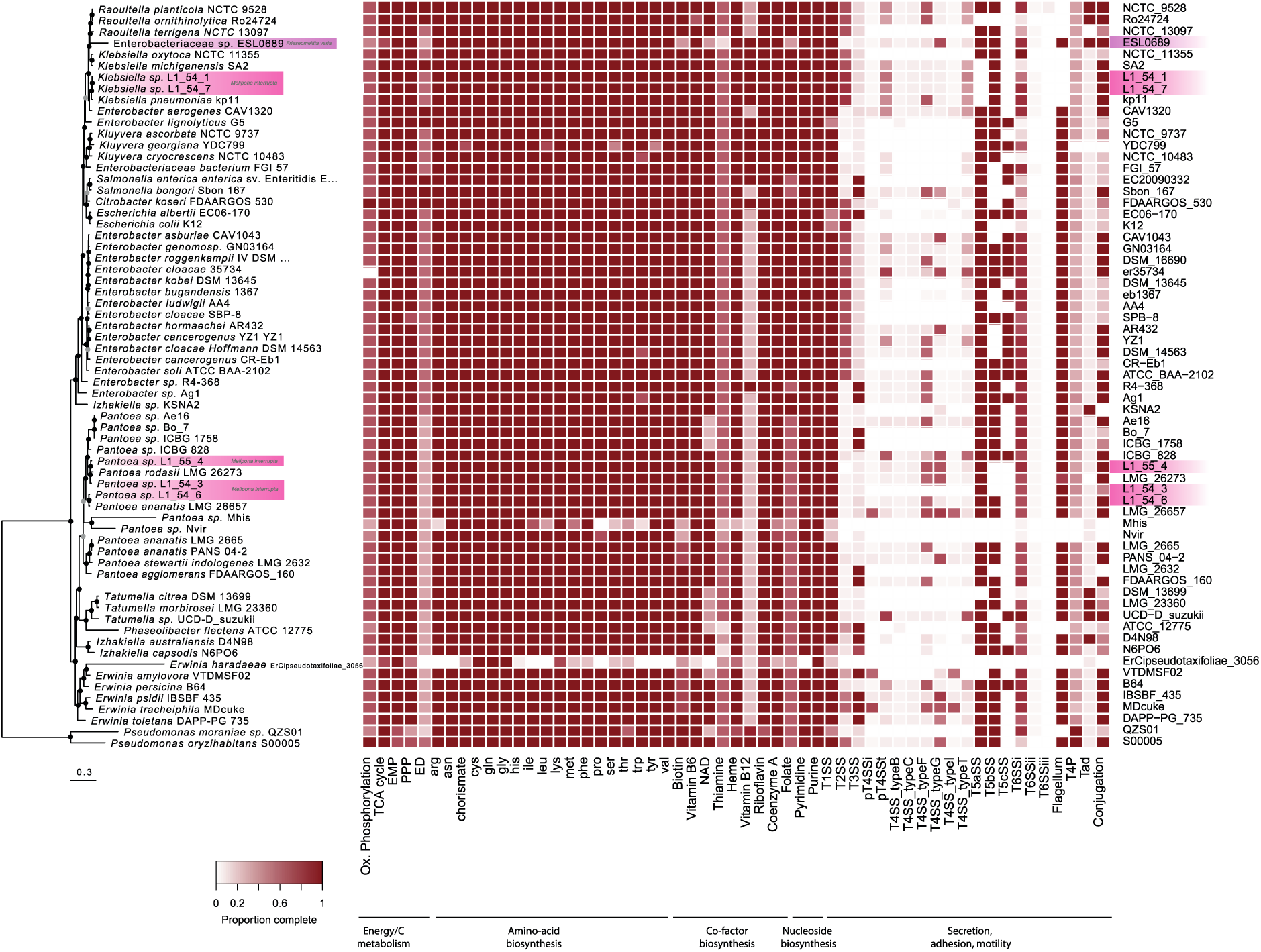
Core genome phylogeny of the Enterobacteriaceae. The maximum likelihood tree was computed on the concatenated amino-acid sequences of 303 single copy core genes using LG+F+I+G4 substitution model. Metabolic heatmap of the Enterobacteriaceae isolates, the genomes are ordered according to the phylogeny. The heatmap indicates the genomic completeness of major metabolic pathways and functions related to energy and carbon metabolism (EMP: Embden–Meyerhof–Parnas), the biosynthesis of amino-acid, co-factors, and nucleoside, as well as secretion, adhesion, and motility across the sequenced bacterial isolates.

**Supplementary Figure 5.**
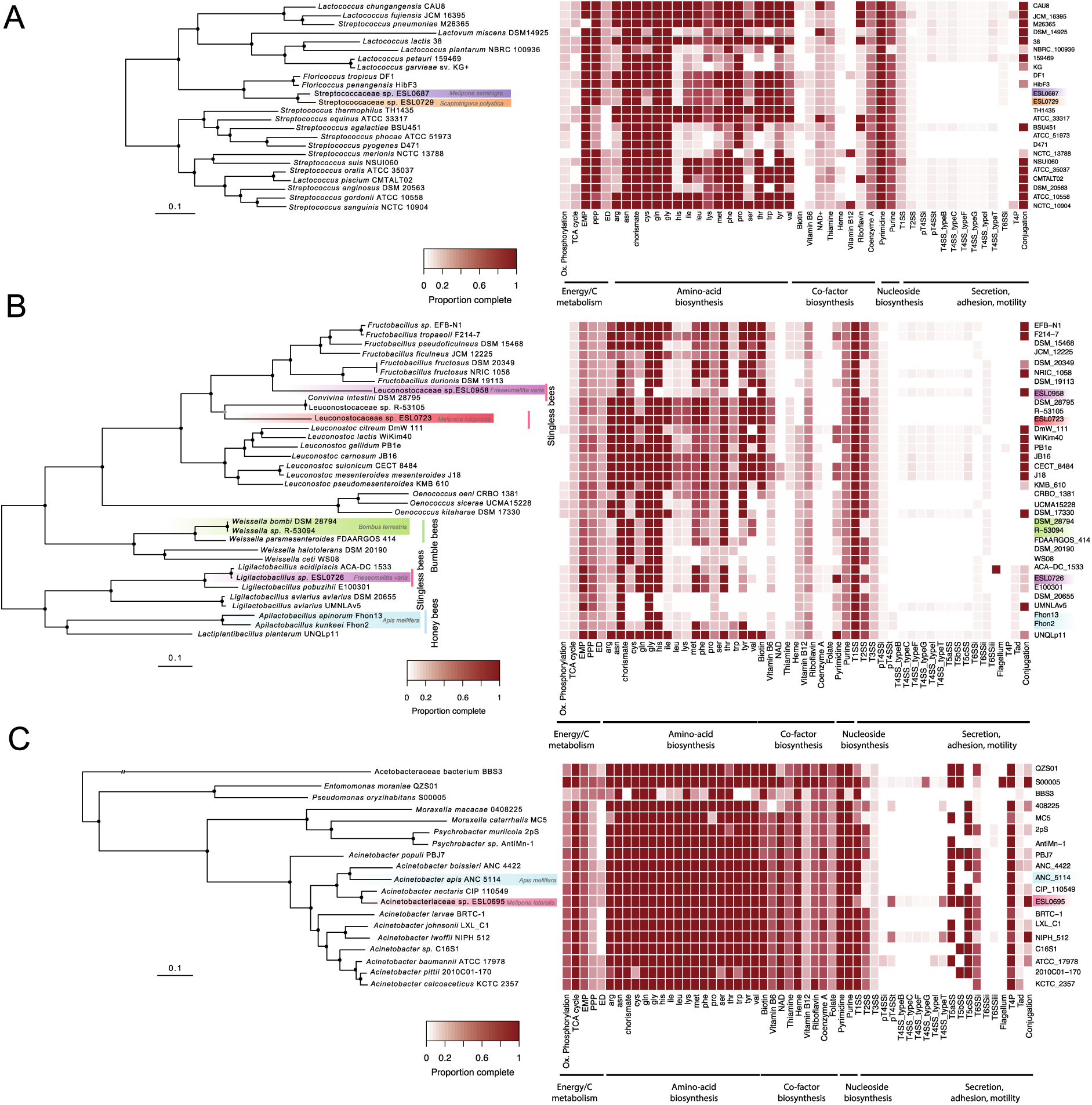
**(A)** Core genome phylogeny of the Streptococcaceae including *Floricoccus* isolates. The maximum likelihood tree was computed on the concatenated amino-acid sequences of 303 single copy core genes using LG+F+I+G4 substitution model. Metabolic heatmap of the Streptococcaceae isolates, the genomes are ordered according to the phylogeny. The heatmap indicates the genomic completeness of major metabolic pathways and functions related to energy and carbon metabolism (EMP: Embden–Meyerhof– Parnas), the biosynthesis of amino-acid, co-factors, and nucleoside, as well as secretion, adhesion, and motility across the sequenced bacterial isolates. **(B)** Core genome phylogeny of the Leuconostocaceae. The maximum likelihood tree was computed on the concatenated amino-acid sequences of 303 single copy core genes using LG+F+I+G4 substitution model. Bee isolates are distributed in two distant monophyletic clades. Metabolic heatmap of the *Leuconostocaceae* isolates, the genomes are ordered according to the phylogeny. The heatmap indicates the genomic completeness of major metabolic pathways and functions related to energy and carbon metabolism (EMP: Embden–Meyerhof–Parnas), the biosynthesis of amino-acid, co-factors, and nucleoside, as well as secretion, adhesion, and motility across the sequenced bacterial isolates. **(C)** Core genome phylogeny of the Moraxellaceae. The maximum likelihood tree was computed on the concatenated amino-acid sequences of 303 single copy core genes using LG+F+I+G4 substitution model. Metabolic heatmap of the Moraxellaceae isolates, the genomes are ordered according to the phylogeny. The heatmap indicates the genomic completeness of major metabolic pathways and functions related to energy and carbon metabolism (EMP: Embden–Meyerhof–Parnas), the biosynthesis of amino-acid, co-factors, and nucleoside, as well as secretion, adhesion, and motility across the sequenced bacterial isolates.

**Supplementary Figure 6.**
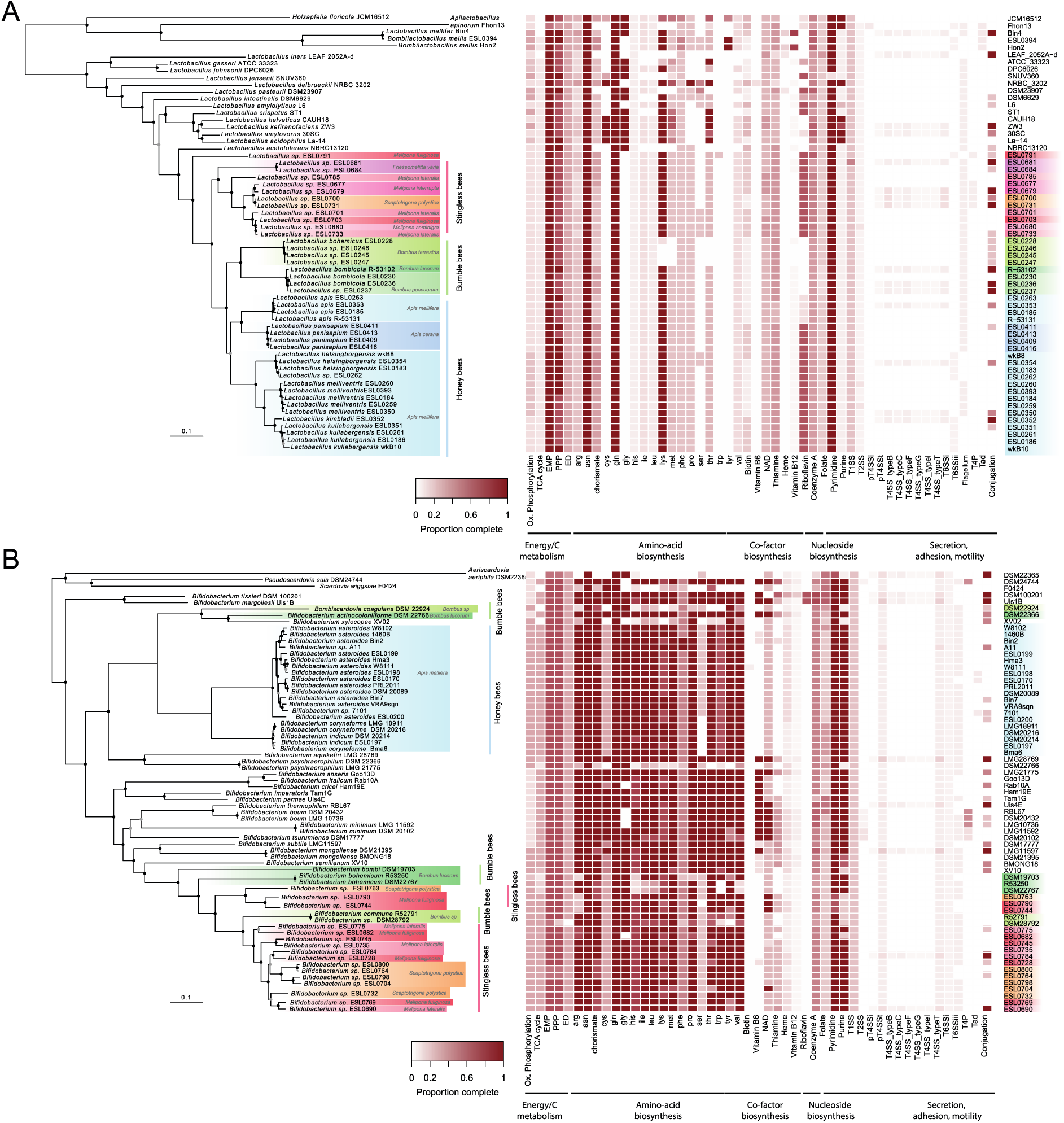
**(A)** Genome-wide phylogeny of the genus *Lactobacillus* including bacteria belonging to the social bee-specific phylotype *Lactobacillus* Firm-5 highlighted in different colors according to the host species/group. The maximum likelihood tree was computed on the concatenated amino acid sequences of 355 single copy core genes using the substitution model LG+F+I+G4. Metabolic heatmap of the *Lactobacillus* isolates, the genomes are ordered according to the phylogeny. The heatmap indicates the genomic completeness of major metabolic pathways and functions related to energy and carbon metabolism (EMP: Embden–Meyerhof–Parnas), the biosynthesis of amino-acid, co-factors, and nucleoside, as well as secretion, adhesion, and motility across the sequenced bacterial isolates. **(B)** Genome-wide phylogeny of the genus *Bifidobacterium* with bacteria belonging to social bee-specific clades. The maximum likelihood tree was computed on the concatenated amino-acid sequences of 151 single copy core genes using the substitution model LG+F+I+G4. Metabolic heatmap of the *Bifidobacterium* isolates, the genomes are ordered according to the phylogeny. The heatmap indicates the genomic completeness of major metabolic pathways and functions related to energy and carbon metabolism (EMP: Embden–Meyerhof–Parnas), the biosynthesis of amino-acid, co-factors, and nucleoside, as well as secretion, adhesion, and motility across the sequenced bacterial isolates.

## Supplementary Tables legends

**Supplementary Table 1.** Gut Homogenates metadata of the 18 stingless bee hives. For each bee species, individuals were collected from three hives in a rural meliponary

**Supplementary Table 2.**

ASV table of our 18 samples. For each ASV, the assigned taxonomy and the number of reads in each sample are displayed. In the sheet RelAbundance, read counts were transformed to relative abundance for each sample. In the Absence_Presence, 1 indicates that at least one read matched the ASV, 0 indicates that the ASV was not found in the sample. Finally, the Family_relAbundaces tab is a summary of the ASVs abundance from each taxonomic family per sample.

**Supplementary Table 3.** ASV table. For each ASV, the assigned taxonomy and the number of reads in each sample are displayed. This analysis groups our 18 samples and samples from W. K. Kwong et al., Dynamic microbiome evolution in social bees. Sci Adv 3, 905 e1600513 (2017). In the sheet RelAbundance, read counts were transformed to relative abundance for each sample. In the Absence_Presence, 1 indicates that at least one read matched the ASV, 0 indicates that the ASV was not found in the sample. Finally, the Family_relAbundaces tab is a summary of the ASVs abundance from each taxonomic family per sample.

**Supplementary Table 4:** Stingless bee strain collection. For each isolate, the bacterial species, bee host are indicated. We blasted the 16S rRNA gene sequences against the NCBI nr database and extracted for each isolate the reference type strain ID and the % of identity between the type strain and our isolate 16S rRHA gene sequences. The Sequenced_Isolates tab, indicates which strains were sequenced.

